# Including variability in air temperature warming scenarios in a lake simulation model highlights uncertainty in predictions of cyanobacteria

**DOI:** 10.1101/734285

**Authors:** Arianna I. Krinos, Kaitlin J. Farrell, Vahid Daneshmand, Kensworth C. Subratie, Renato J. Figueiredo, Cayelan C. Carey

**Affiliations:** Department of Biological Sciences, Virginia Tech, Blacksburg, Virginia USA; Department of Electrical and Computer Engineering, University of Florida, Gainesville, Florida USA; Odum School of Ecology, University of Georgia, Athens, Georgia USA

**Keywords:** climate change, blooms, phytoplankton, GLM, Mendota, probability distributions

## Abstract

Despite growing evidence that climate change will increase temperature variability and the frequency of temperature extremes, many modeling studies that analyze the effects of warming scenarios on cyanobacteria in lakes examine uniform warming temperature scenarios without including any variability. Here, we used the one-dimensional hydrodynamic General Lake Model coupled to Aquatic EcoDynamics modules (GLM-AED) to simulate 11 years of nitrogen-fixing and non-nitrogen-fixing cyanobacterial biomass in Lake Mendota (Madison, WI, USA). We developed climate scenarios with either uniform (constant) warming or variable warming based on random sampling of daily air temperatures from either a normal or Poisson probability distribution. We found that while the median cyanobacterial biomass among repeated simulations for each of the years was similar regardless of whether or not air temperature variability was included in the climate scenarios, the randomly-sampled air temperature distribution scenarios exhibited much greater variability in the year-to-year cyanobacterial biomass estimates. Our results suggest that including temperature variability in climate scenarios may substantially change our understanding of how climate warming may alter cyanobacterial blooms over yearly to decadal time scales. To more effectively predict the range of possible future cyanobacterial dynamics, both the magnitude and variability of warming must be considered when developing climate scenarios for lake modeling studies.

## 1. Introduction

Freshwater lakes are responding globally to changing air temperatures (Chapra et al., 2017; IPCC, 2014; Michalak, 2016; O’Reilly et al., 2015; Richardson et al., 2017; Scheffers et al., 2016; Wood et al., 2016). In addition to increasing overall mean air temperatures, anthropogenic greenhouse gas emissions are increasing the variability of daily and seasonal temperatures (Cox et al., 2018; Fischer and Schär, 2009; Otto-Bliesner et al., 2016; Räisänen and Räisänen, 2002; Trenberth et al., 2015). The magnitude and seasonality of predicted temperature changes vary substantially among geographic regions and temporal scales (IPCC, 2014). In the north temperate latitudes of North America, warming is projected to be greater in winter than in summer, with a substantial increase in extreme summertime heat events (Romero-Lankao et al., 2014). The effects of increased air temperature variability on freshwater ecosystems could potentially exceed that of increases in mean air temperature (Settele et al., 2014), and thus it is important that warming scenarios take temperature variability into account (Dillon et al., 2016).

To date, most lake modeling studies have either applied a constant degree of warming in their temperature scenarios (e.g., a uniform -1 to +4□ added every day to the model driver data; e.g., Elliott, 2010; Elliott et al., 2016) or derived warming projections from a suite of climate models (Bruce et al., 2018; Elliott et al., 2006; Li et al., 2016), though neither of these approaches accounts for day-to-day variability in air temperatures. While some lake modeling studies have used multiple temperature scenarios, most have not incorporated within-scenario variability (e.g., Butcher et al., 2015). Focusing on uniform temperature scenarios, defined here as applying a daily increase in temperature in a constant way to all days in the simulation, excludes the increased variability in air temperatures that is predicted to occur due to climate change (IPCC, 2014; Romero-Lankao et al., 2014).

Increasing mean air temperatures and temperature variability will likely play a substantial role in shaping lake phytoplankton communities (e.g., Carey et al., 2012; Havens, 2008; Paerl and Huisman, 2008; Visser et al., 2016a). Phytoplankton are responsible for much of the primary production in lakes, and are an important part of freshwater food webs (Creed et al., 2018; Stockner and Porter, 1988; Vogt et al., 2017; Zwart et al., 2015). Different phytoplankton taxa have different temperature sensitivities (reviewed by Reynolds 2006), which will influence their growth rate responses to future temperature warming and its variability. High growth rates of phytoplankton can result in blooms, which both degrade the aesthetic value of freshwater ecosystems and contribute to toxic scums (Havens, 2008; Paerl et al., 2016). In particular, air temperature warming may increase the magnitude and frequency of cyanobacterial blooms (Carey et al., 2012; Chapra et al., 2017; Harris et al., 2016; Paerl, 2016; Paerl and Huisman, 2008; Visser et al., 2016). Cyanobacteria have higher thermal optima than eukaryotic phytoplankton (up to 35°C for most taxa; Reynolds 2006), so higher temperature variability could increase the incidence of very warm days on which the optimum temperature for cyanobacteria occurs in the lake (Konopka and Brock, 1978; Visser et al., 2016; Walls et al., 2018; Walter Helbling et al., 2015). However, like most lake modeling studies, most work to date studying the effects of warming on phytoplankton has focused on changes to mean temperatures, not temperature variability (e.g., De Stasio et al., 1996; Mooij et al., 2005; Striebel et al., 2016; Thomas and Litchman, 2015), despite evidence that variability in water temperatures can alter phytoplankton (Edlund et al., 2017; Havens et al., 2016; Sahoo et al., 2016; Saros et al., 2016; Woolway and Merchant, 2017, 2018).

Here, we used a lake ecosystem simulation model to answer the question, How does increased mean temperature and daily temperature variability affect the biomass of lake cyanobacteria? We calibrated a one-dimensional hydrodynamic water quality model to simulate two phytoplankton functional groups that represent the physiological and ecological traits of nitrogen-fixing (N-fixing) and non-nitrogen-fixing (non-N-fixing) cyanobacteria, and quantified the effects of warming and more variable air temperatures on phytoplankton community dynamics. We developed scenarios of variable air temperature forcing by randomly sampling from different distributions of air temperature increases on each day of the simulation. We expected that including air temperature variability would result in an increased incidence of cyanobacterial blooms, because especially warm days might trigger cyanobacterial blooms. We also expected that the results would vary substantially among the multiple years in a simulation due to background variability in other meteorological drivers (e.g., precipitation), which were not modified in the air temperature scenarios.

## 2. Methods

### 2.1 Study site description

Lake Mendota is a large, dimictic, eutrophic lake in Madison, Wisconsin, USA (Table 1). Mendota has a history of summer (e.g., May - October) phytoplankton blooms that are dominated by cyanobacteria and lead to degraded water quality and beach closures (Beversdorf et al., 2013; Brock, 1985; Weirich et al., 2019). The lake has been monitored for several decades as part of the North Temperate Lakes Long-Term Ecological Research (NTL-LTER; lter.limnology.wisc.edu) program and the Global Lakes Ecological Observatory Network (GLEON; gleon.org). Routine depth profiles of water temperature, concentrations of dissolved oxygen (DO), nutrients (nitrogen [N] and phosphorus [P]), and total filtered chlorophyll-*a*, and phytoplankton grab samples are manually collected at the deepest point in the lake approximately every two weeks during the ice-free period (roughly April - October) and a few times during the ice-covered period. High-frequency sensors have been deployed each year during the ice-free period since 2006, and provide sub-daily water temperature profiles and DO concentrations at 0.5 m depth.

**Table 1.**
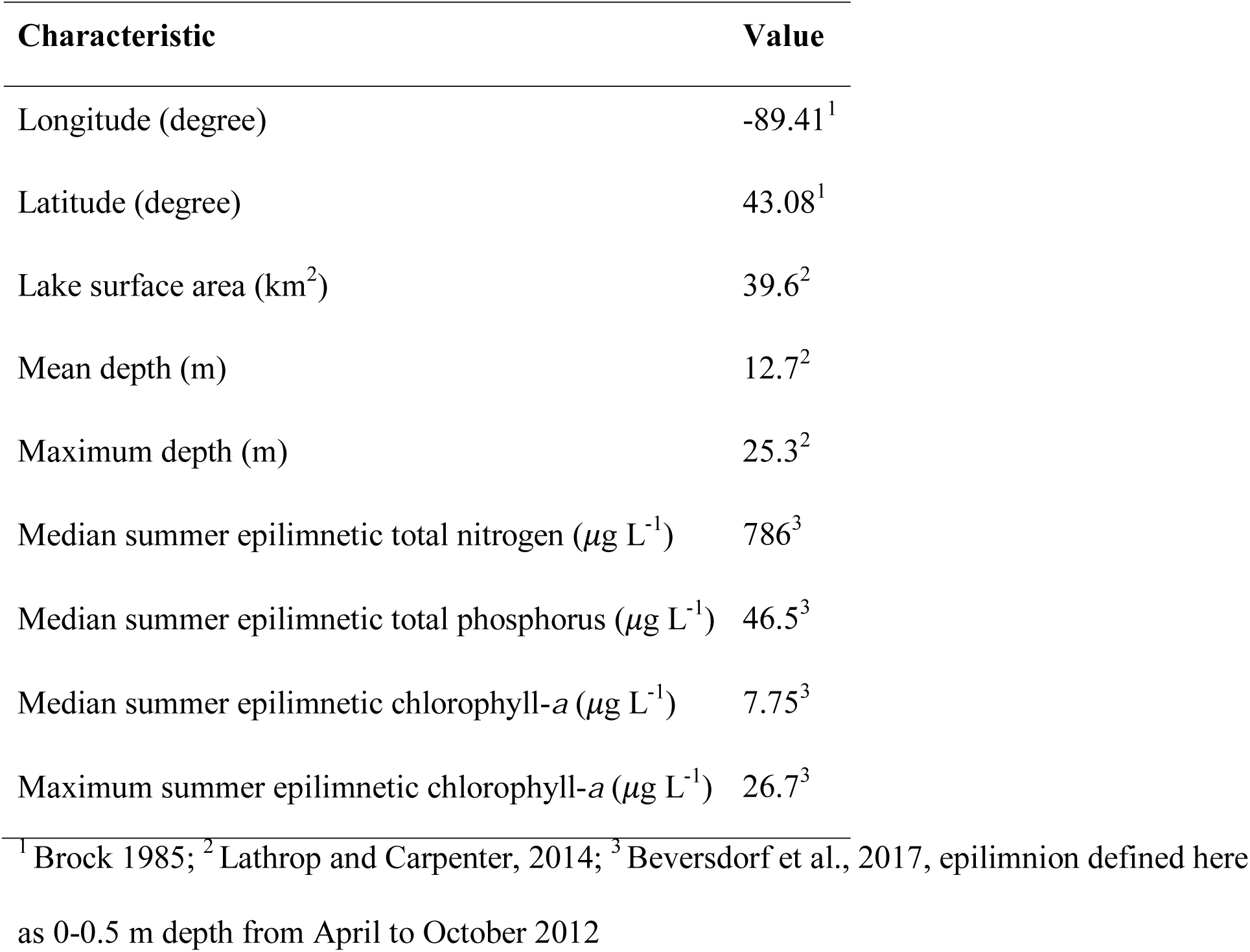
Physical, chemical, and biological characteristics of Lake Mendota, Madison, Wisconsin, USA.

| Characteristic | Value |
| --- | --- |
| Longitude (degree) | -89.41<sup>1</sup> |
| Latitude (degree) | 43.08<sup>1</sup> |
| Lake surface area (km<sup>2</sup>) | 39.6<sup>2</sup> |
| Mean depth (m) | 12.7<sup>2</sup> |
| Maximum depth (m) | 25.3<sup>2</sup> |
| Median summer epilimnetic total nitrogen ($\mu\text{g L}^{-1}$) | 786<sup>3</sup> |
| Median summer epilimnetic total phosphorus ($\mu\text{g L}^{-1}$) | 46.5<sup>3</sup> |
| Median summer epilimnetic chlorophyll-*a* ($\mu\text{g L}^{-1}$) | 7.75<sup>3</sup> |
| Maximum summer epilimnetic chlorophyll-*a* ($\mu\text{g L}^{-1}$) | 26.7<sup>3</sup> |

<sup>1</sup> Brock 1985; <sup>2</sup> Lathrop and Carpenter, 2014; <sup>3</sup> Beversdorf et al., 2017, epilimnion defined here as 0-0.5 m depth from April to October 2012

### 2.2 Model setup & observational data

We used the one-dimensional hydrodynamic General Lake Model (GLM v. 2.1.8; Hipsey et al., 2014, 2019) coupled with the Aquatic EcoDynamics (AED) library (Hipsey et al., 2013; henceforth GLM-AED) to explore potential effects of air temperature warming on cyanobacterial blooms in Lake Mendota. GLM simulates lake water and energy budgets, while AED simulates dynamic water quality, including concentrations of nutrients (e.g., N, P) and phytoplankton functional groups (e.g., non-nitrogen-fixing and nitrogen-fixing cyanobacteria). We chose this model in part because meteorological drivers (e.g., air temperatures) can be easily modified to simulate a variety of climate change scenarios. In addition, the model’s low computational requirements made it ideally suited for running a multi-year model simulation many times. We ran GLM-AED over an 11-year period from 8 November 2003 to 31 December 2014 at an hourly time step. This time period contains the most complete time series of observational data available and includes representative climatic events for Mendota, including both flood and drought years (e.g., 2010 and 2009, respectively; see Usinowicz et al., 2016).

The GLM-AED uses meteorological and surface water inflow and outflow driver data to produce a dynamic lake water and energy profile (Hipsey et al., 2019). We compiled hourly meteorological driver data from the full simulation period (2003-2014), including air temperature (°C), wind speed (m s^−1^), relative humidity (%), short and longwave radiation (W m^−2^), and precipitation (m d^−1^), from the North American Land Data Assimilation System (NLDAS-2).

Our model simulated three inflows to Mendota, two of which were based on observed data from streamflow gauges at the Yahara River at Highway 113 (United States Geological Survey [USGS] ID #05427850) and Pheasant Branch (USGS ID #05427948). The third inflow (“Balance”) was estimated as the difference between the gauged outflow (USGS ID #05428500) volume and the two gauged inflows, and represented contributions to the water budget from runoff, groundwater, and smaller tributaries. Gaps in daily flow data were filled using linear interpolation, and all flow rates were transformed from USGS units of ft^3^ s^−1^ to m^3^ s^−1^. Daily surface inflow driver data were based on a combination of these observed and statistically-modeled flow rates (m^3^ s^−1^), water temperature (°C), and inorganic and organic fractions of N, P, and carbon (all mmol m^−3^; described below). Inflow water temperatures were based on available gauge measurements. Missing water temperature data were estimated as a function of observed air temperature using breakpoint regression between measured mean daily air temperatures and measured water temperatures at the Yahara Highway gauge. Inflow water temperatures did not vary among air temperature scenarios. Outflow data included flow rate from the USGS gauge for the Yahara River outflow.

Daily inflow nutrient concentrations were based on data collated from USGS gauges on Pheasant Branch and the Yahara River at Windsor (USGS ID #05427718). Nutrient concentrations from the Yahara River were applied to the “Balance” inflow, as it is likely more representative of ungauged portion of the Mendota watershed (Carpenter et al., 2018; Lathrop, 1992). Inflow driver data included concentrations of eight different P and N fractions. Phosphorus fractions included filterable reactive P (FRP), adsorbed FRP (FRP that is chemically bound to particles), dissolved organic P (DOP), and particulate organic P (POP). Nitrogen inflows included nitrate (NIT), ammonium (AMM), dissolved organic N (DON), and particulate organic N (PON).

We used USGS data for total P (TP; USGS parameter #00665) collected at both the Yahara Windsor and Pheasant gauges to estimate concentrations of individual P fractions. We estimated adsorbed FRP as equal to gauged TP (P.C. Hanson, pers. comm.). We estimated FRP as 46.97% of measured TP, based on the calculated median proportion of TP as measured orthophosphate (which we assumed to be functionally equivalent to FRP) between 1 January 2011 and 31 December 2015. We estimated DOP as 39.00% of measured TP based on Rigler (1964)’s measurements of DOP as a proportion of TP for a 0.45 µm filter pore size used by the USGS. We approximated POP by subtracting the DOP and FRP fractions from 100%, resulting in POP as 14.03% of TP. All P concentrations were transformed from mg L^−1^ to mmol m^−3^ using the molar mass of P (30.97).

Because N data were not collected at any of the focal USGS gauges prior to 1 October 2010, we estimated daily concentrations of each N fraction for each gauge (Yahara Windsor, Pheasant) using a linear regression between gauged flow volume and available measurements of NIT and AMM. We estimated PON concentrations by subtraction based on measured AMM plus organic N (USGS parameter #00625) and AMM alone (USGS parameter #00608) for each gauge. DON concentrations were set to zero. All N concentrations were transformed from mg L^−1^ to mmol m^−3^ using the molar mass of N (14.01).

Particulate organic carbon (POC) and dissolved organic carbon (DOC) were included in inflow data as a static median concentration for all inflows using data collected from Lake Mendota tributaries between 15 April 2016 and 14 November 2016 (Hart et al., 2017).

The AED phytoplankton module was configured with four phytoplankton functional groups: N-fixing and non-N-fixing cyanobacteria, diatoms, and chlorophytes, which account for the majority of the phytoplankton biomass in Mendota (Snortheim et al., 2017). The N-fixing cyanobacterial functional group approximates the traits of phytoplankton in the genera *Aphanizomenon* and *Dolichospermum* (formerly *Anabaena*), while the non-N-fixing cyanobacteria are modeled after phytoplankton in the genus *Microcystis* (Kara et al., 2012). Phytoplankton biomass in GLM-AED is simulated as a function of mortality, respiration, nutrient uptake, and vertical movement. Zooplankton, which have rates associated with grazing, respiration, predation, other mortality, and metabolism, graze upon phytoplankton (Hipsey et al., 2013, 2019). We used only parameters available in the default configuration of GLM-AED v.2.1.8 (see Table A1).

### 2.3 Model calibration & validation

Initial calibration of the baseline GLM-AED model was based on previous GLM models set up for Mendota (Kara et al., 2012; Snortheim et al., 2017). Calibration focused on capturing seasonal trends in epilimnetic (0-2 m) water temperature, DO concentrations, and the timing and peak biomass for N-fixing and non-N-fixing cyanobacteria. Model outputs were compared to NTL-LTER high-frequency buoy data and manual samples for water temperature and DO, and manual samples of phytoplankton cell counts and bulk chlorophyll-*a* concentrations (Magnuson et al., 2010a, b, c, d). Biomass of both N-fixing and non-N-fixing cyanobacteria was estimated from phytoplankton count data following Eppley et al. (1970), Menden-Deuer and Lessard (2000), and Strathmann (1967).

Calibration involved sequential manual tuning of phytoplankton parameters following the approach in Kara et al. (2012), with the goal of minimizing root mean square error (RMSE; Table 2) of summer (June – September) epilimnetic (0-2 m) total chlorophyll-*a* (μg L^−1^) and the biomass (mmol C m^−3^) of N-fixing and non-N-fixing cyanobacteria at 1 m throughout the year (Table 2). Calibration focused on parameters related to growth and respiration, including optimum temperature, growth rate, and respiration-associated biomass losses (see Table A1). We focused on epilimnetic (0-2 m depth) concentrations because this depth is where the majority of cyanobacterial blooms occur (Weirich et al., 2019).

**Table 2.**
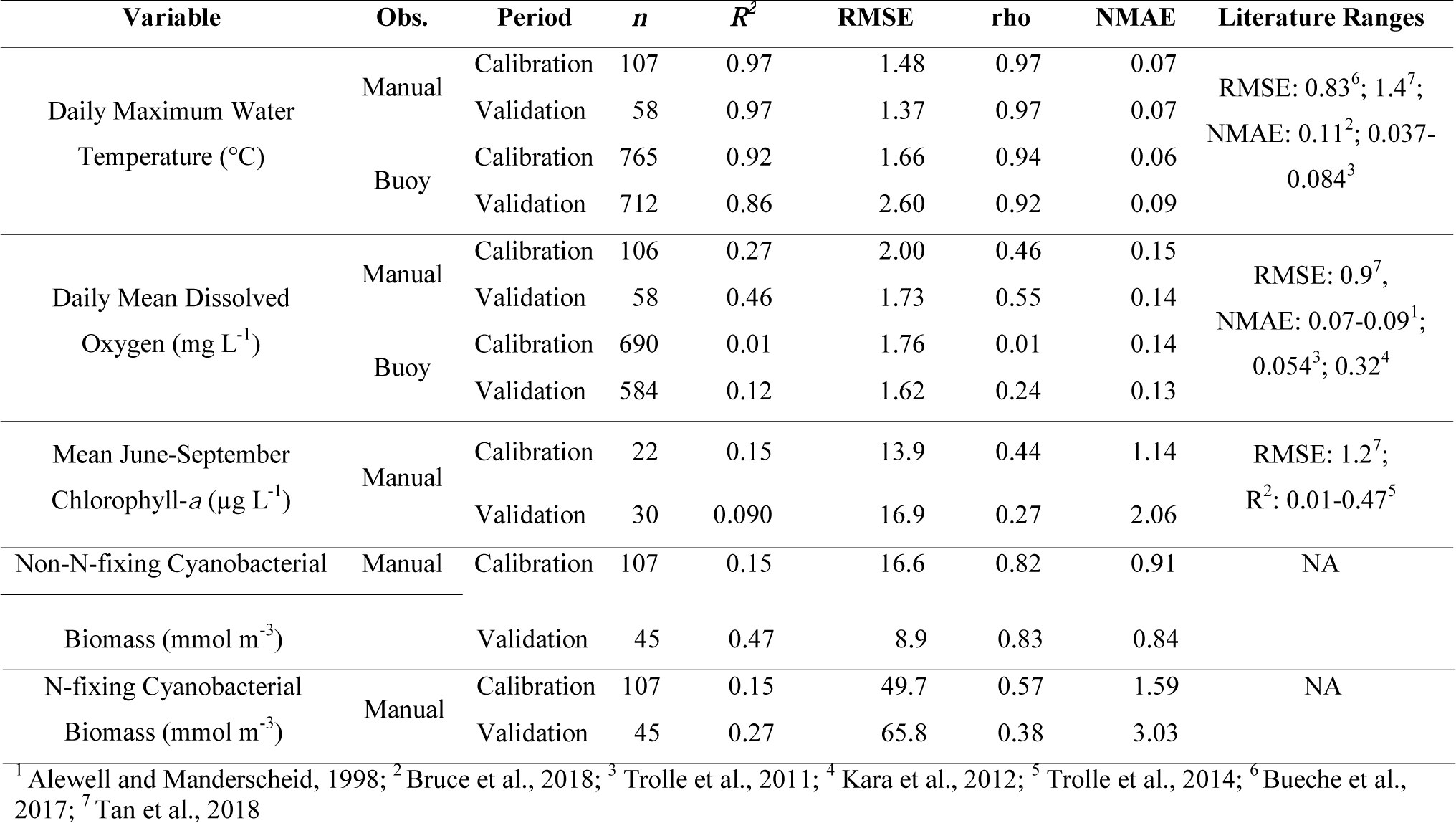
Goodness-of-fit (GOF) metrics (coefficient of determination [R^2^], root mean square error [RMSE], Spearman’s rank correlation coefficient [rho], and normalized mean absolute error [NMAE]) for focal state variables comparing the baseline calibrated model with observed data. GOF metrics are reported separately for the calibration (1 January 2004 – 31 December 2010) and validation (1 January 2011 – 31 December 2014) periods. All GOF metrics were calculated for the lake surface (0-1 m, water temperature and manual dissolved oxygen; 0.5 m, buoy dissolved oxygen; 0-2 m, manual chlorophyll-*α*and N-fixing and non-N-fixing cyanobacterial biomass) based on available manual and buoy data. Literature ranges indicate the range of GOF reported for other studies of Lake Mendota.

**Table 3.** Median yearly biomass (in mmol C m^−3^) for each temperature distribution scenario and mean temperature offset for (a) non-N-fixing cyanobacteria and (b) N-fixing cyanobacteria. Values for non-uniform distributions are the median across repeated replicates (n=10 replicates per distribution). Among the temperature distribution scenarios, the median values aggregated over the 10-year simulation did not substantially vary.

| (a) **Distribution** | +0°C | +1°C | +2°C | +3°C | +4°C | +5°C |
| --- | --- | --- | --- | --- | --- | --- |
| Uniform | 7.91 | 14.21 | 9.72 | 8.22 | 9.40 | 7.73 |
| Normal | 7.71 | 10.78 | 11.61 | 9.44 | 9.52 | 8.73 |
| Tail 2 | 7.91 | 11.29 | 11.21 | 11.09 | 10.54 | 9.04 |
| Tail 4 | 7.91 | 11.98 | 11.34 | 10.49 | 9.67 | 8.18 |
| Tail 8 | 7.91 | 11.62 | 11.75 | 10.15 | 10.40 | 8.37 |
| (b) **Distribution** | +0°C | +1°C | +2°C | +3°C | +4°C | +5°C |
| Uniform | 21.98 | 19.77 | 24.53 | 25.97 | 25.15 | 25.69 |
| Normal | 23.23 | 19.75 | 22.05 | 22.32 | 24.55 | 23.18 |
| Tail 2 | 21.98 | 20.93 | 21.96 | 22.57 | 22.89 | 23.16 |
| Tail 4 | 21.98 | 21.41 | 22.54 | 22.91 | 21.55 | 23.49 |
| Tail 8 | 21.98 | 21.26 | 20.36 | 20.75 | 21.33 | 22.71 |

We calculated multiple goodness-of-fit metrics (Table 2) including RMSE, the coefficient of determination (R^2^), Spearman’s rho, and normalized mean absolute error (NMAE; Alewell and Manderscheid, 1998) separately for calibration (1 January 2004 - 31 December 2010) and validation (1 January 2011 - 31 December 2014) periods.

### 2.4 Climate scenarios

We combined air temperature offsets and probability distributions to assess the effects of both the magnitude and variability of climate warming on phytoplankton. Simulated air temperature offsets (from +0 to +5°C) applied to the GLM-AED meteorological driver file for the modeling period (2003-2014) encompassed a range of potential warming scenarios, as in Subratie et al., (2017). Climate projections were based on MACAvs-METDATA downscaled global climate models (Abatzoglou, 2013; Abatzoglou and Brown, 2012; Taylor et al., 2012), which project air temperature warming of approximately +5.4°C above historical conditions (1950-1980) for Madison, Wisconsin by 2099 under RCP8.5.

We incorporated variability into the climate warming scenarios by randomly sampling three different probability distributions (uniform, normal, or Poisson) to define the mean and standard deviation of daily air temperature offsets applied throughout the model period. First, the uniform scenarios applied a static, integer increase in daily air temperature relative to observed air temperatures. The uniform scenarios ranged in integer offsets from +0°C to +5°C, and did not include any air temperature variability. Second, air temperature offsets were generated by sampling from a symmetric normal distribution centered around the mean of each integer air temperature offset (+0°C to +5°C) with a standard deviation of 1°C (Figure 1). For each day of the model simulation, the air temperature offset was randomly drawn from the normal distribution. For normal distribution scenarios with a mean increase of +0°C, daily air temperature offsets could be cooler (up to -1°C) than the observed baseline temperature. Third, we generated air temperature offset scenarios for three Poisson distributions. Warming scenarios based on Poisson distributions had a mean normalized to be within 0.1°C of each uniform air temperature offset from +0 to +5°C and a lambda (λ) equal to 2, 4, or 8 (“Tail 2”, “Tail 4”, “Tail 8”, hereafter). These values of lambda were chosen because they represent a range of spread of the Poisson distribution (Figure 1), and thus encompassed a range of variability for day-to-day air temperature offsets. For the normal distribution and each of the Poisson distributions (the “temperature distribution scenarios”), we ran 10 replicate model simulations to capture a range of potential effects driven by day-to-day variability in the air temperature offset within the distribution. Because the uniform scenarios applied the same air temperature offset each day and the GLM-AED model is deterministic, no replicate uniform simulations were run.

**Figure 1.**
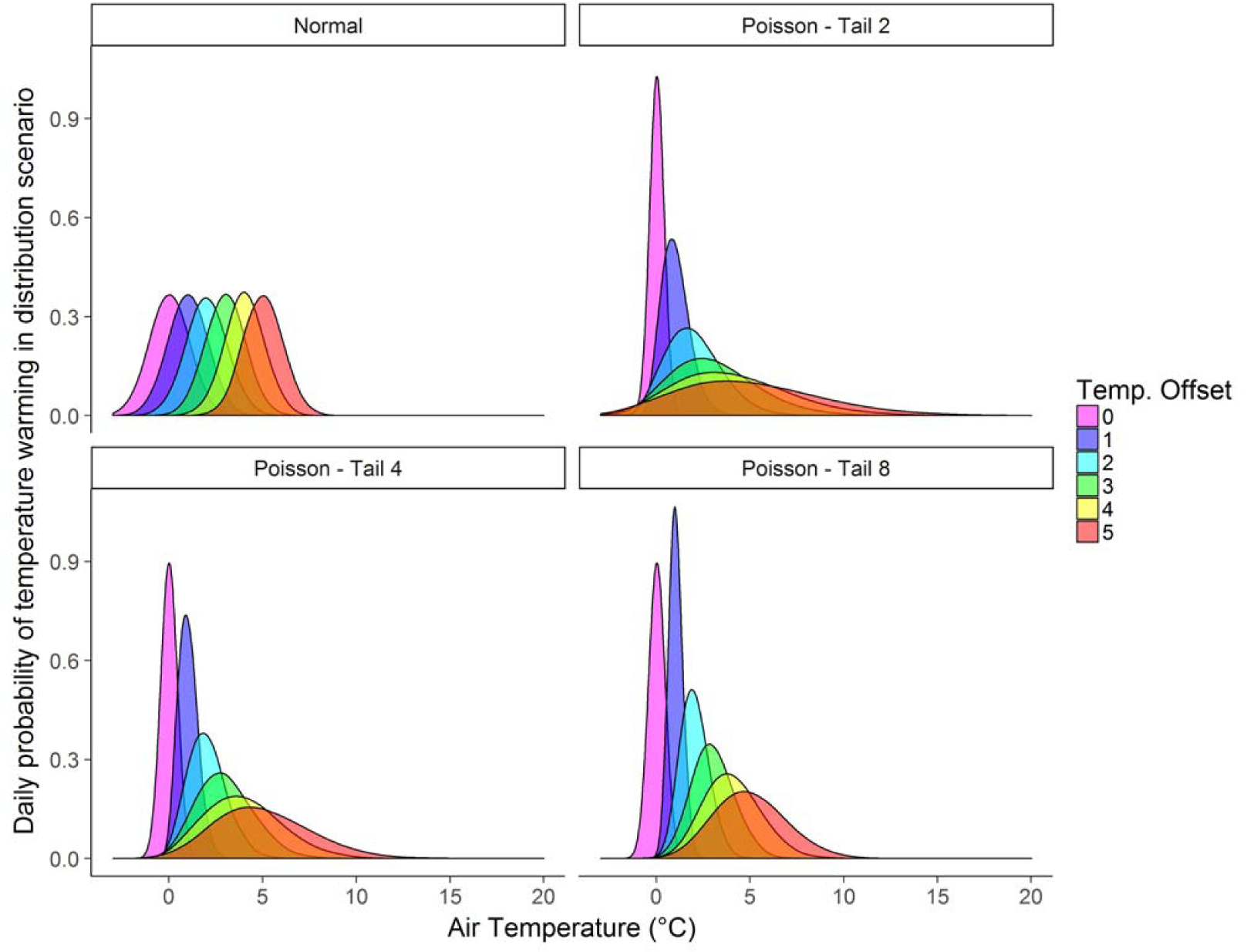
Distributions from which daily air temperature changes were applied to the 11-year simulation period for the non-uniform temperature distribution scenarios (Normal, Poisson -Tail 2, Poisson - Tail 4, and Poisson - Tail 8). Colors indicate the mean air temperature offset for each temperature distribution scenario (+0°C to +5°C).

### 2.5 Bloom quantification

We assessed the effects of the magnitude and variability of climate warming on cyanobacterial blooms using two metrics: (1) the number of days per year that surpassed a bloom threshold and (2) the maximum yearly cyanobacterial biomass. We also calculated the overall median biomass across the 10-year simulation period and all scenario replicates. Metrics were calculated separately for N-fixing and non-N-fixing cyanobacteria, and years were based on calendar year (January 1 – December 31). For each metric, we estimated yearly values as the median or maximum value among replicates for a given air temperature offset (e.g., +1, +2, +3°C) and distribution. We used 5 µg L^−1^ (29.2 mmol C m^−3^) of chlorophyll-*a* as the threshold for a cyanobacterial bloom, following Boyer et al. (2009). While this concentration is lower than the chlorophyll-*a* criterion for lakes and reservoirs set by many states (US Environmental Protection Agency, 2019), our study focused on two individual cyanobacterial functional groups, rather than the full phytoplankton community. We converted modeled daily cyanobacterial carbon concentrations (mmol C m^−3^) to chlorophyll-*a* concentrations using the molar mass of C (12.01) and the carbon to chlorophyll-*a* ratio (estimated as 70 for Mendota phytoplankton; P.C. Hanson, pers. comm).

### 2.6 Statistical analyses

For each of the two bloom metrics (number of days per year exceeding the bloom threshold and maximum yearly biomass) and cyanobacterial groups, we used Anderson-Darling tests (Engmann and Cousineau, 2011; Scholz and Stephens, 1987) to quantify differences between the distribution of model outputs (henceforth, “output distribution”) for each air temperature offset (+0°C to +5°C). Statistical significance was based on Bonferroni-corrected α for multiple comparisons. We evaluated model output from 1 January 2005 to 31 December 2014, excluding 2003 and 2004 as a spin-up period, for a total of n=10 simulation years.

Each output distribution was composed of 10 points representing annual values for each of 10 model simulation years (2005-2014). We compared the bloom metrics from the uniform warming scenarios to the modeled temperature distribution scenarios or the baseline scenario (+0°C), rather than to the observational data. For the normal and Poisson distributions, we calculated a single bloom metric for each model simulation year aggregated across the 10 scenario replicates. We examined whether the most extreme number of bloom days per year and maximum biomass per year were significantly different from either the baseline scenario (+0°C) or the uniform distribution for each temperature offset. By comparing the warming scenario model output to the baseline model output, not observational data, our analyses examined the temperature responses of an idealized eutrophic lake based on Mendota. All analyses were conducted using R 3.5.3 (R Core Team, 2019).

## 3. Results

### 3.1 Calibration results

The GLM-AED model provided a reasonable fit to the observational data, with our calculated R^2^ and RMSE goodness-of-fit metric values within the range reported by similar previous lake numerical simulation models for temperature, dissolved oxygen, and chlorophyll (Bruce et al., 2018; Kara et al., 2012; Snortheim et al., 2017; Trolle et al., 2011, 2014), and we observed overall good correspondence between the observed and modeled seasonal dynamics and peak biomass for both cyanobacterial groups (Figure 2). The model R^2^ was 0.47 and 0.27 for non-N-fixing cyanobacteria and N-fixing cyanobacteria, respectively, during the validation period (Table 2), indicating reasonable correspondence between the observed and modeled data.

**Figure 2.**
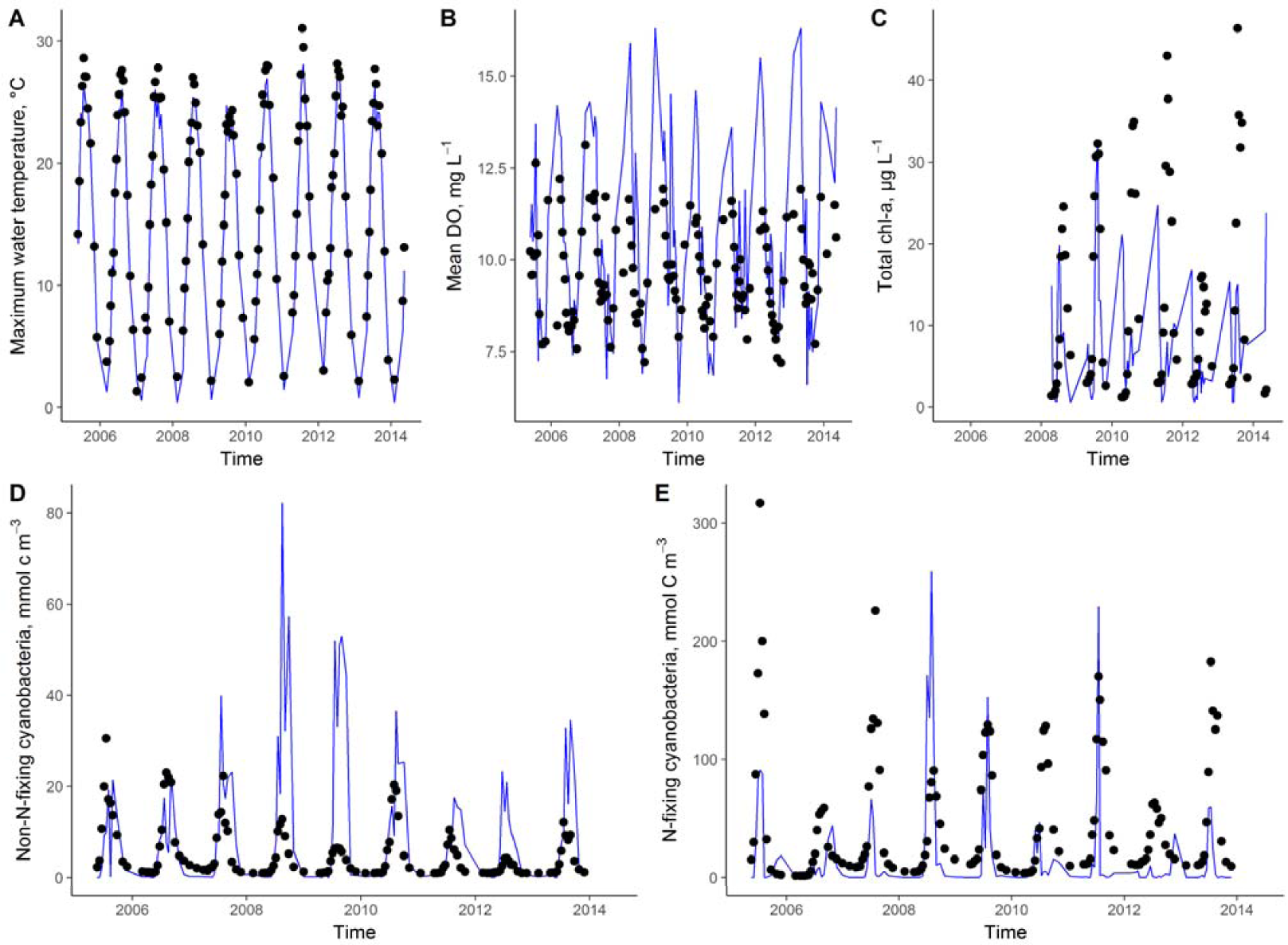
Baseline modeled (blue lines) versus observed (black points) for manually-collected observations at Lake Mendota’s deepest site for daily epilimnetic A: maximum water temperature (°C; 0-1 m), B: mean dissolved oxygen (DO, mg L^−1^; 0-1 m), C: June-August chlorophyll-*a* (µg L^−1^; 0-2 m), D: non-N-fixing cyanobacteria (mmol C m^−3^; 0-2 m), and E: N-fixing cyanobacteria (mmol C m^−3^; 0-2 m).

We observed a much stronger correlation between the observed and modeled data for non-N-fixing cyanobacteria than N-fixing cyanobacteria. For example, the validation RMSE for non-N-fixing cyanobacteria was 8.9 mmol C m^−3^, while the RMSE for N-fixing cyanobacteria was 65.8 mmol C m^−3^ (Table 2). Similarly, the Spearman’s rho was 0.84 for non-N-fixing cyanobacteria versus 0.38 for N-fixing cyanobacteria during the validation period and the NMAE was 0.84 and 3.03 for non-N-fixing and N-fixing cyanobacteria, respectively. Goodness-of-fit metrics were generally much better for the model in the validation period than the calibration period, likely because it was shorter in duration.

### 3.2 Effect of air temperature on surface water temperature

Epilimnetic lake water temperatures (1 m depth) varied with both mean air temperature offset and the temperature distribution scenario used (Tables B1-4; Figure 3). Among all scenarios, water temperatures warmed less than the mean air temperature offset: in the uniform scenario, the mean summer water temperature increase across the 10-year simulation ranged from 0.39 (+1°C air temperature offset) to 1.56°C (+5°C temperature offset) (Figure 3). Maximum daily summer water temperatures were significantly warmer than the baseline for the +1 to +5°C air temperature offsets (Table B1), and median daily water temperatures were also significantly different from the baseline for all +1 to +5°C temperature offsets (Table B2).

**Figure 3.**
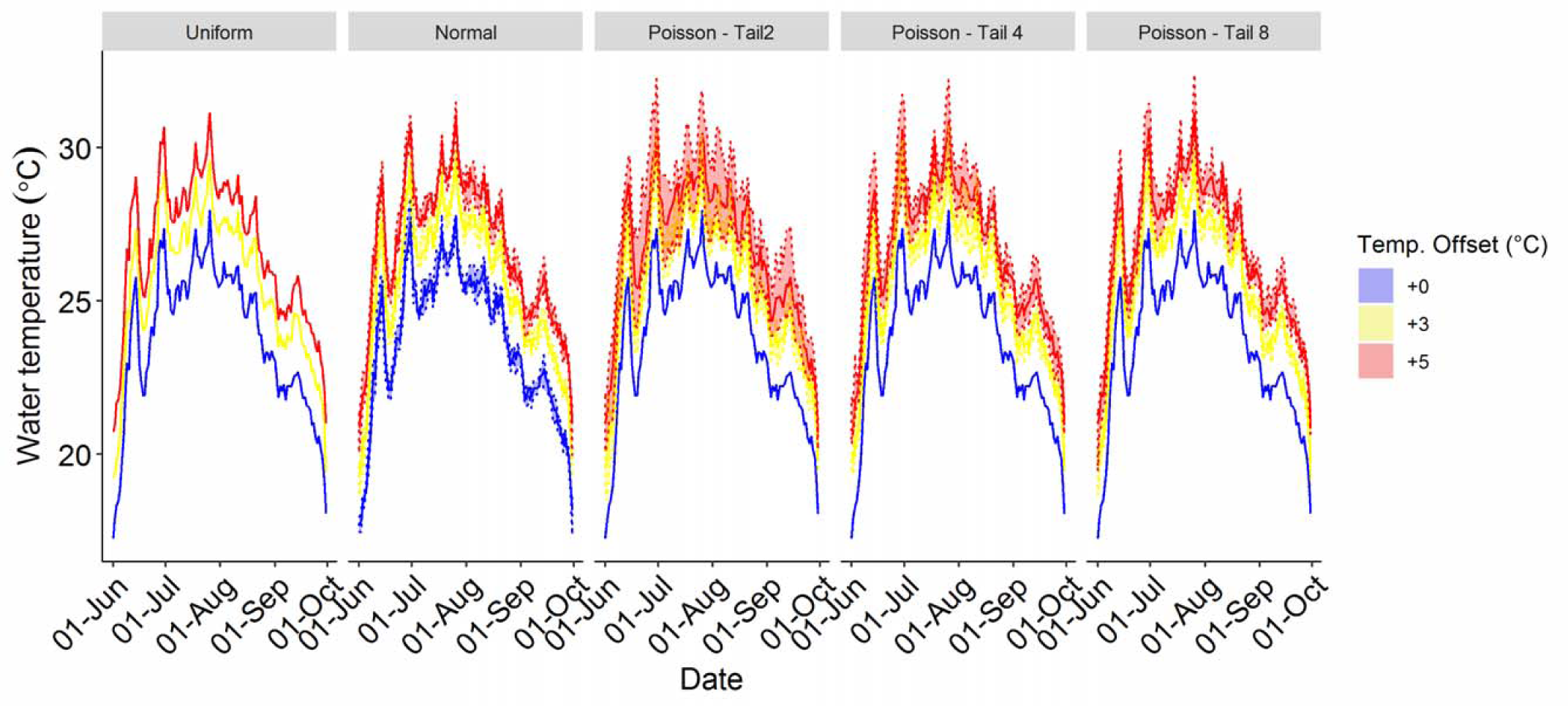
Modeled daily surface (1 m) water temperatures (°C) in Lake Mendota under +0°C, +3°C, and +5°C mean air temperature offsets for each temperature distribution scenario (Uniform, Normal, Poisson Tail 2, Poisson Tail 4, and Poisson Tail 8), shown from 1 June to 1 October 2005 as an example year. Solid lines indicate daily median water temperatures; dashed lines for non-uniform distributions indicate daily minimum and maximum water temperatures among replicates (n=10 per distribution).

Daily water temperatures varied among replicates in the non-uniform temperature distribution scenarios (Figure 3), but the distributions of median daily epilimnetic water temperatures across replicates at each temperature offset were not significantly different from the uniform scenario (Table B3). However, the distributions of maximum daily epilimnetic water temperatures were significantly different from the uniform scenario for the +2 to +5°C temperature offsets for the normal and all Poisson distributions, one Poisson distribution was significant at +1°C, and the normal distribution was always significantly different from the uniform distribution (Table B4), with much more variability in day-to-day water temperatures in the non-uniform scenarios.

### 3.3 Effect of distribution on number of days exceeding the bloom threshold each year

Overall, we found that the median number of bloom days for both non-N-fixing and N-fixing cyanobacteria increased with air temperature warming among all temperature distribution scenarios. In the uniform warming scenarios for non-N-fixing cyanobacteria, the median number of days per year surpassing our defined bloom threshold increased by 289% for our lowest temperature offset, +1°C, from 18 days per year (year-to-year range: 0-85 days) in the baseline simulation to 70 days per year (year-to-year range: 16-109) for +1°C; by 211% to 56 days per year (year-to-year range: 0-109) for +4°C warming; and by 119% to 40 days per year (year-to-year range: 0-67) for +5°C of warming (Figure 4A).

**Figure 4.**
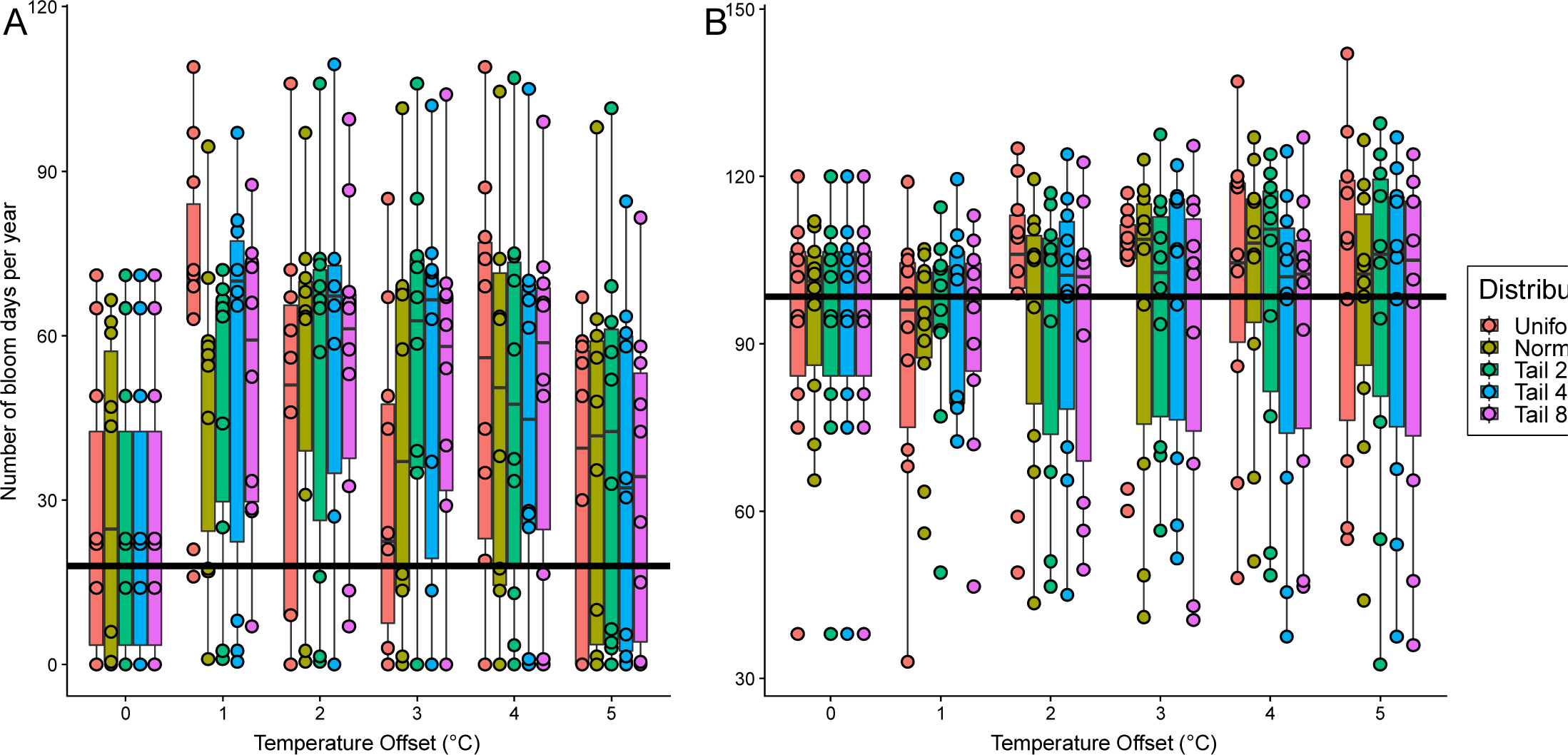
Days per year that exceeded the 5 µg L^−1^ chlorophyll-*a* bloom threshold for A: non-N-fixing cyanobacteria and B: N-fixing cyanobacteria as a function of mean temperature offset (+0°C to +5°C). Boxplots represent different temperature distribution scenarios across temperature offsets; points within each boxplot represent a year within the model simulation (2005-2014). For non-uniform temperature distribution scenarios, yearly points are medians among replicates (n = 10). Horizontal line indicates the median bloom days per year in the baseline scenario (+0°C) among all model years (2005-2014). Note that the extent of y-axes differs between panels.

The median number of non-N-fixing bloom days per year also increased for the simulations with non-uniform distributions of warming temperatures, with the greatest increases in the median number of bloom days occurring in the “Tail 2” Poisson distribution (increased by 26 days at 4°C; 142% increase) and the lowest increase in the median number of bloom days occurring in the “Tail 4” Poisson distribution (increased by 11 days at 4°C; 58% increase). The “Tail 8” Poisson and normal distributions were intermediate, with an increase of 13 bloom days (72% increase) and 25 days (125% increase) at 4°C, respectively, for non-N-fixing cyanobacteria. However, the median yearly number of bloom days in the uniform scenario or any of the temperature distribution scenarios (with the exception of +1°C for the uniform distribution; Table B5) were not significantly different from the baseline, nor were the temperature distribution scenarios significantly different from the uniform distribution (Table B6).

While the model output indicated that non-N-fixing cyanobacteria increased overall with warming air temperature despite not being significant, variability in cyanobacterial responses, as assessed by the among-year range in the number of bloom days, was higher in the normal and Poisson temperature distribution scenarios than in the uniform warming scenario (Figure 5). While the among-year range in bloom days for the uniform distribution was 93 days (16 to 109 days per year) in the +1°C air temperature offset, the among-year range was 1.3 times higher (123 days) in the normal and Poisson distributions (0 to 123 days per year). For the +5°C temperature offset, the range for the normal and Poisson distributions was nearly two times higher than the uniform distribution (123 days, 0 to 123 days per year and 67 days, 0 to 67 days per year, respectively). A maximum range of 0 to 103 bloom days was observed for a single year and temperature offset among replicates (2009, +3°C temperature offset; Figure 5).

**Figure 5.**
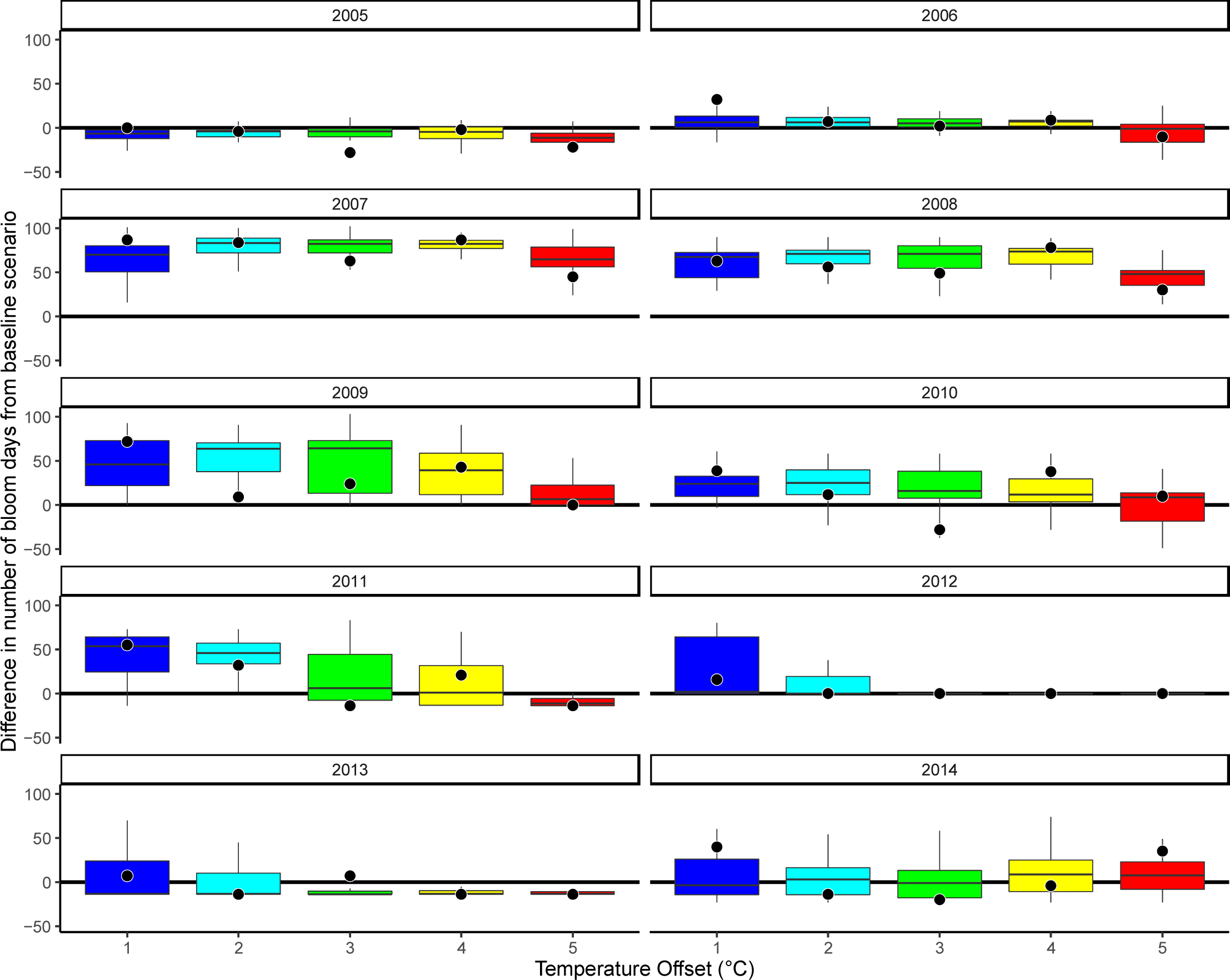
Number of bloom days for each modeled year (2005-2014) for non-N-fixing cyanobacteria. The filled black circles indicate the number of bloom days for the uniform distribution scenario and the horizontal line indicates the number of bloom days in the baseline scenario (+0°C). The boxplots encompass the number of bloom days among all four non-uniform distribution scenarios and their 10 replicates (n=40 total) across the range of mean temperature offsets (+1°C - +5°C). Among years, the difference between the effect of the uniform distribution scenario and the non-uniform distributions scenarios on the number of bloom days was variable.

We observed similar patterns in increasing median number of bloom days per year for N-fixing cyanobacteria in warming simulations (Figure 4B), despite this pattern not being significant (Table B7 and B8). At 5°C warming, the median number of bloom days per year in the uniform distribution scenario increased by 10% from 98.5 with no warming (year-to-year range: 38-120) to 108.5 days (year-to-year range: 18-144). Thus, while N-fixing cyanobacteria were predicted to increase overall, individual years exhibited higher, lower, or the same number of bloom days compared to the baseline scenario (Figure 4B). The Tail 2, 4, and 8 Poisson distributions had similar increases in the median number of N-fixing cyanobacterial bloom days per year (6-7%), increasing to approximately 105 bloom days in the +5°C warming scenario compared to 98.5 in the baseline (+0°C) scenario. The normal distribution exhibited a smaller increase (2%) in the number of N-fixing cyanobacterial bloom days per year, from a median of 102 (+0°C) to 104 (+5°C) bloom days. a range of 0 to 145 bloom days among all years and temperature offsets and up to 124 bloom days for a single year and temperature offset was observed for N-fixing cyanobacteria (2014, +3°C temperature offset; Figure 6), while the range of bloom days across all years and temperature offsets for the uniform distribution was 33 to 142 bloom days.

**Figure 6.**
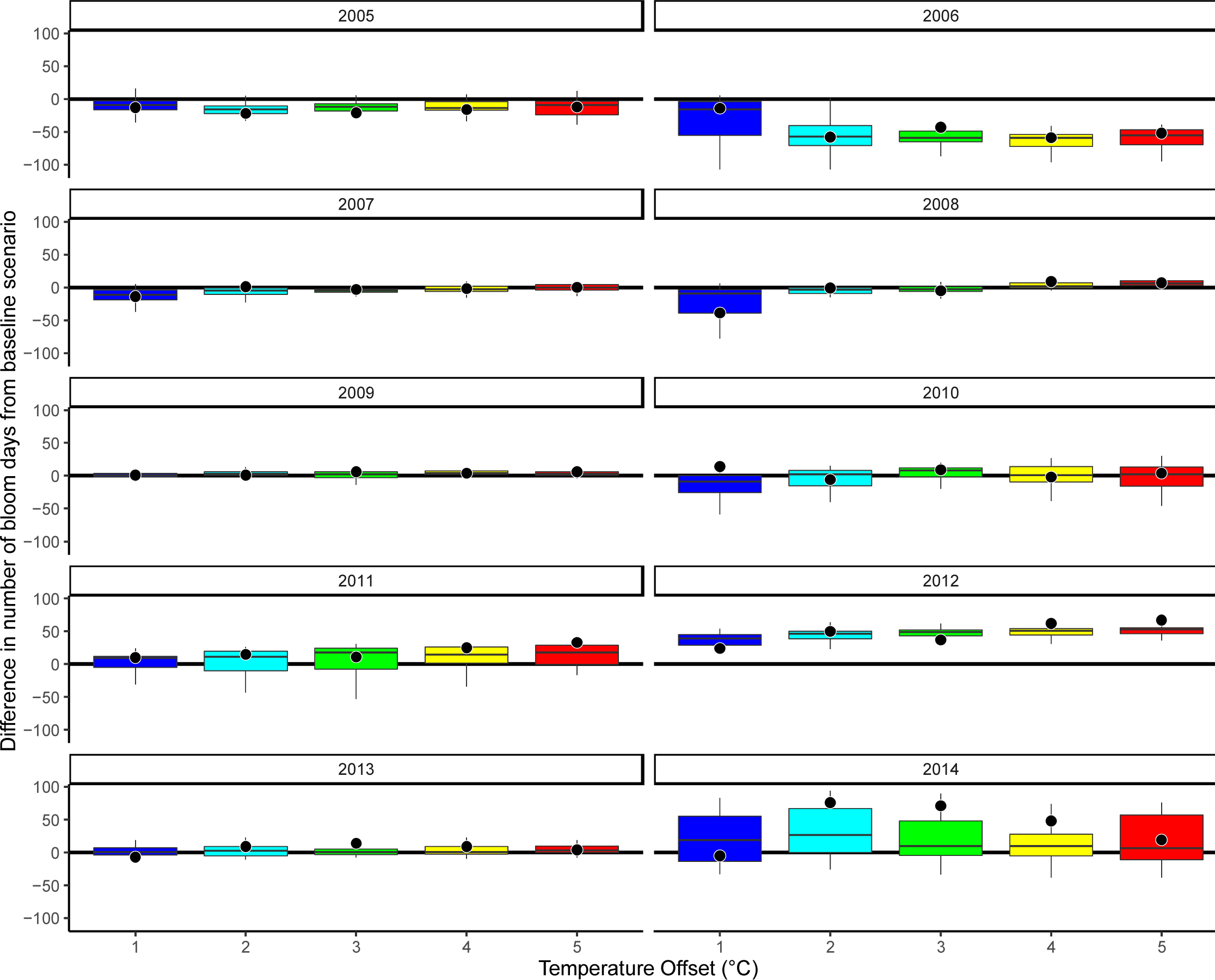
Number of bloom days for each modeled year (2005-2014) for N-fixing cyanobacteria. The filled black circles indicate the number of bloom days for the uniform distribution scenario and the horizontal line indicates the number of bloom days in the baseline scenario (+0°C). The boxplots encompass the number of bloom days among all four non-uniform distribution scenarios and their 10 replicates (n=40 total) across the range of mean temperature offsets (+1°C - +5°C). Among years, the difference between the effect of the uniform distribution scenario and the non-uniform distributions scenarios on the number of bloom days was variable.

While the median number of bloom days per year were generally not significantly different between non-uniform and uniform distributions among temperature offsets for either non-N-fixing or N-fixing cyanobacteria (Tables B5 and B7, the maximum number of bloom days per year exhibited more statistically significant differences among distributions, though not at all temperature offsets (Tables B9 and B10). The greatest bloom duration in each of the non-uniform temperature distribution scenarios was significantly different from the uniform case for at least one of the temperature offsets from +1 to +5°C, though not for all of the temperature offsets or for both cyanobacterial groups (Tables B9 and B10). For N-fixing cyanobacteria, these differences were most pronounced at +1°C warming and disappeared at the +4-5°C temperature offsets (Table B9).

For non-N-fixing cyanobacteria, most temperature distribution scenarios were significantly different from the baseline at the +2°C and +3°C temperature offsets (Table B10). At higher levels of warming (3-5°C), more non-N-fixing cyanobacteria number of bloom day values were statistically significantly different than N-fixing cyanobacteria values, with 6 out of 12 significant results for non-N-fixing cyanobacteria versus 2 out of 12 significant results for N-fixing cyanobacteria (Tables B9 and B10). For non-N-fixing cyanobacteria maximum bloom days per year, the uniform distribution was only significantly different from the baseline scenario for the +1°C temperature offset, while most temperature offsets were significantly different for the non-uniform distributions, with the exception of the +5°C temperature offset for the Poisson Tail 4 and Tail 8 scenarios (Table B11). For N-fixing cyanobacteria, the uniform distribution was never significantly different from the baseline, while all four non-uniform distributions were significant or marginally significant at +1-2°C of warming, 3 at +3°C, 2 at +4°C, and 3 at +5°C (Table B12).

### 3.4 Effect of distribution on peak yearly biomass

Similar to median number of bloom days for both N-fixing and non-N-fixing groups, maximum yearly biomass exhibited notable year-to-year differences among temperature distribution scenarios, but the among-year median of the maximum yearly biomass was not significantly different among temperature distributions, other than for the Poisson distributions at +5°C for non-N-fixing cyanobacterial (Figure 7; Tables B13, B14) nor across temperature offsets within a distribution (Tables B15, B16). However, for both cyanobacterial functional groups, we observed a large range in the year-to-year model output of biomass. Among both uniform and non-uniform distributions, there was substantial variability among temperature distribution replicates. For example, the Tail 2 and Tail 4 distributions at 4 and 5°C exhibited maximum yearly biomass concentrations for non-N-fixing cyanobacteria that were lower than in the baseline scenario (up to 44.5 mmol m^−3^ lower biomass than in the baseline). The output distribution composed of the overall yearly maximum biomass among the 10 replicates for each temperature distribution exhibited at least one statistically significant difference from both the baseline scenario and the uniform temperature distribution for the nonzero temperature offsets (Tables B17 to B20). While there were consistent statistically significant differences between the non-uniform temperature distributions and the baseline scenario for non-N-fixing cyanobacteria for between +1°C and +4°C temperature offsets (Table 17), these differences were often marginally or not significant for N-fixing cyanobacteria (Table 19).

**Figure 7.**
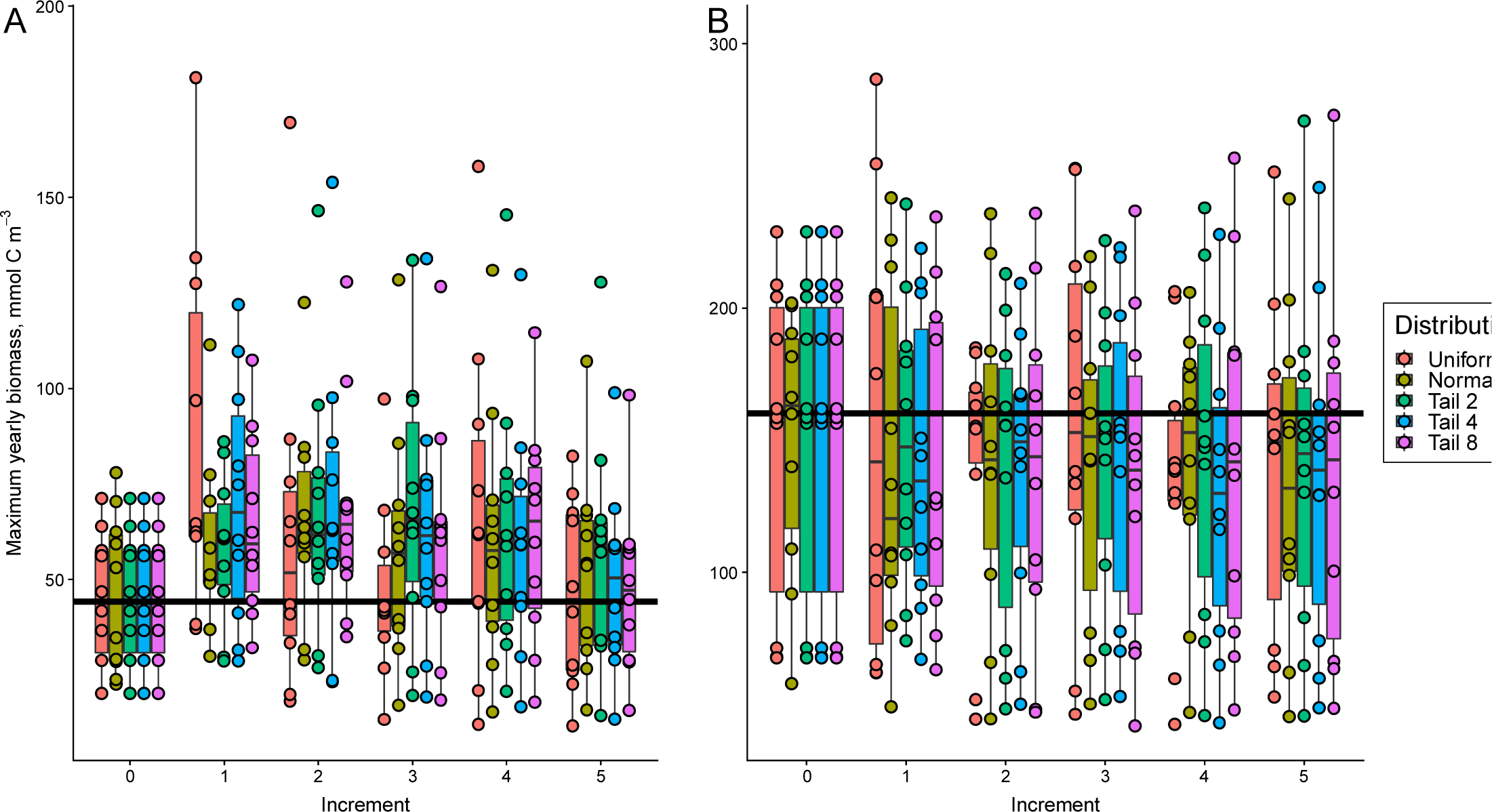
Maximum yearly biomass (mmol C m^−3^) of A: non-N-fixing cyanobacteria and B: N-fixing cyanobacteria as a function of mean temperature offset (+0°C to +5°C). Boxplots represent different temperature distribution scenarios across temperature offsets; points within each boxplot represent the maximum biomass for each year within the model simulation (2005-2014), and the median among years is indicated by the horizontal line within each boxplot. For non-uniform temperature distribution scenarios, yearly points are medians of the maxima among replicates (n = 10). Horizontal black line indicates the median maximum biomass across simulation years in the baseline scenario (+0°C) among all model years (2005-2014). Note that the extent of y-axes differs between panels.

## 4. Discussion

Our results suggest that incorporating temperature variability into warming scenarios for simulation modeling may influence confidence in the magnitude and direction of cyanobacterial bloom predictions. Randomly sampling air temperature distributions to generate warming scenarios did not always reproduce the cyanobacterial biomass model output generated by uniform warming scenarios, even if the overall mean air temperature increased by the same offset, because the maximum biomass predicted was significantly different than the uniform scenario for several of these distributions (Tables B11, B12, B18, and B20). Our analysis showed that the projected number of bloom days per year and peak biomass concentrations for cyanobacteria have roughly the same mean and median after aggregating across 10 years and repeated random simulations, but that there was a difference up to 100 bloom days per year and up to 300 mmol C m^3^ maximum yearly biomass (median difference 128.4 mmol C m^3^ for N-fixing cyanobacteria; 53.9 mmol C m^3^ for non-N-fixing cyanobacteria) among individual temperature distribution scenario replicates. In some cases, the impact of warming in the median case was greatest at low levels of warming (+1 to +2°C temperature offset), and was rarely significant at high levels of warming (+4 to +5°C; e.g., Table B20). This may be because increased warming increases zooplankton grazing, nutrient limitation, or other factors that constrain cyanobacterial growth.

Thus, including a reasonable amount of day-to-day air temperature variation when applying a warming scenario to a lake model can moderate or even reverse the direction of cyanobacterial responses in comparison to a baseline (non-warming) simulation. In some studies, this could mean a decrease (or no change) in blooms with warming, in contrast to the expectation of greater bloom incidence in a warmer future (e.g., Elliott, 2012). The difference in the modeled number of bloom days per year among the 10 replicates of the same non-uniform distribution ranged up to 100 bloom days within the same year, which has large implications for predictions of future blooms. A range of 100 bloom days per year (∼3 months) represents approximately half of the thermally stratified summer period from May to October for Lake Mendota. Given that temperature variability will increase in the future due to climate change (IPCC, 2014; Romero-Lankao et al., 2014), temperature scenarios for bloom modeling should incorporate that variability, even if it means that bloom predictions will become more uncertain. The number of scenario replicates is thus important; using too few replicates may not capture the expected median, and more replicates could reveal an even greater range in predictions. Including air temperature variability may be especially important when modeling smaller lakes, which have relatively low thermal inertia compared to larger lakes, oceans, and seas (Piccolroaz et al., 2015; Stainsby et al., 2011; Ye et al., 2019), and consequently may be more sensitive to variability in warming temperatures.

We observed that our bloom metrics measured in individual years were less consistent among years under non-uniform warming scenarios (Figures 5 and 6. This is likely because water temperatures could vary substantially depending on the pattern of warming obtained via random daily draws from the chosen distribution (Figure 3), and the maximum daily values of water temperature varied significantly for the normal and Poisson distributions at the +2 to +5°C temperature offsets (Table B4), thereby altering the timing of and frequency with which cyanobacterial thermal optima are met. As a consequence of the process of random drawing from a distribution, water temperatures in individual years may differ, on average, at any given temperature offset. For example, for the Poisson Tail 8 distribution, the median summer water temperature was 1.92°C warmer than the baseline in 2010 but 1.49°C warmer than the baseline in 2011 for the +3°C temperature offset. For the uniform distribution, the median summer water temperature was 3.14°C warmer than the baseline in 2010 and 2.66°C warmer in 2011 for the +5°C temperature offset. We found that both inter-annual and inter-scenario variability in terms of the maximum cyanobacterial biomass and number of yearly bloom days increased when variable temperature warming distribution regimes were applied, relative to the uniform warming scenario.

While non-N-fixing cyanobacteria showed a stronger warming response, particularly at low levels of warming (+1-2°C temperature offsets), N-fixing cyanobacteria exhibited a much greater range of responses among replicates in the randomly-sampled temperature distribution scenarios than non-N-fixing cyanobacteria. The greater warming response of non-N-fixing cyanobacteria at low temperature offsets may be due to the higher optimum temperature of the non-N-fixing cyanobacteria (over 20°C and up to 41°C in comparison to optima below 20°C for many other phytoplankton; Carey et al., 2012; Jöhnk et al., 2008; Lürling et al., 2013; Reynolds, 2006), meaning that the group responded more quickly and strongly to warming water temperatures, as was parameterized in our GLM-AED model (Table A1). The greater variability in N-fixing cyanobacteria summer biomass relative to non-N-fixing cyanobacteria may be because of the higher overall biomass concentrations of this functional group (Figure 2) and the fact that the biomass concentrations were much more poorly represented in our model for N-fixing cyanobacteria as compared to non-N-fixing cyanobacteria. Further, the fact that we were unable to properly capture the dynamics of this group might have been the result of the sensitivity of the group to slight changes in environmental conditions, as parameterized in our model.

The wide range of biomass and bloom days for the two groups of cyanobacteria among individual years and repeated model runs may be caused in part by the fact that greater variability in air temperatures may reduce lake ice cover (Magee et al., 2016). Reduction of ice cover alters thermal stratification and light availability, potentially altering the lake’s mixing regime (Woolway and Merchant, 2019) and level of primary productivity (O’Beirne et al., 2017). These changes are often more pronounced in spring in the Northern Hemisphere due to warmer winter temperatures (Duguay et al., 2006; Sharma et al., 2019). Daily temperature variation, particularly in winter, may also drive decreases in average lake water level (Yongliang et al., 2007; Yuan et al., 2015), which could increase nutrient concentrations (Jeppesen et al., 2015), thereby indirectly altering cyanobacterial biomass.

Our investigation is limited in that we only evaluated two broad cyanobacterial functional trait groups. While this functional group approach is common in many studies (e.g., De Senerpont Domis et al., 2007, 2012; Elliott, 2012; Mooij et al., 2005), it excludes variation that exists among taxa within the same group that might influence responses to warming (Bohnenberger et al., 2018; Cirés and Ballot, 2016; Edwards et al., 2012; Litchman et al., 2010; Litchman and Klausmeier, 2008; Özkundakci et al., 2016; Shan et al., 2019). In addition, we imposed air temperature variation randomly, rather than taking into account potential seasonal patterns expected under climate change, such as greater warming in the winter in the Northern Hemisphere (Hayhoe et al., 2010; IPCC, 2013; Pryor et al., 2013). Using a non-deterministic model would also improve our understanding of how temperature variability in future scenarios alters cyanobacterial responses by self-adjusting to make small changes in air temperatures between repeated trials. This would require more trials or more caveats about the results to account for added uncontrolled variability in the model, but would eliminate the bias involved in choosing a specific distribution for errors. Importantly, while our study only addressed variability in air temperature, other meteorological and land use drivers, including nutrient inputs and precipitation, will likely further increase uncertainty in bloom predictions.

This study provides evidence that the more variable the weather is from day to day, the less certainty there is in the magnitude and direction of future change in cyanobacterial blooms. While we found that the median cyanobacterial biomass and bloom days aggregated over 10 simulation years were approximately the same among all warming offsets and temperature distribution scenarios, the maximum and minimum year-to-year predictions were very different. Our analysis highlights that the range or uncertainty in cyanobacterial responses to warming cannot be evaluated from a single deterministic simulation. This could be addressed by treating every simulation study like a comparison of models of different climate change trajectories, such as is being done by groups like the Inter-Sectoral Impact Model Comparison Project (ISIMIP; isimip.org). While we only evaluated the effects of air temperature in this study, our results suggest that other sources of meteorological variability could also be important, and we recommend including additional meteorological drivers such as precipitation and wind speed in future modeling work, which will likely be important for studying the variability of cyanobacterial blooms under different warming scenarios.

Finally, this study provides evidence that temperature variability may be as important a driver of change in ecosystems as warming itself. Future studies should investigate how the frequency, magnitude, and timing of air temperature changes affect the variability of bloom responses. For example, if different levels of warming occur at different times of year (following Hayhoe et al., 2010), does that increase or decrease the predictability of blooms throughout the year? Although our results are limited to a single lake model experiment, we demonstrate that exploring different patterns of air temperature warming and variability is needed to inform cyanobacterial bloom modeling in the face of climate variability and change.

## Acknowledgments

This work was supported by the U.S. National Science Foundation [grant numbers EF-1702506, DEB-1753639, ICER-1517823, CNS-1737424, ACI-1234983]. We thank Paul C. Hanson for guidance in early model development, and members of the Carey Lab for helpful feedback. Observational data used in this project were obtained from the North Temperate Lakes-LTER program.

## Author Contribution Statement

CCC, RJF, AIK, and KJF conceived ideas underlying the project; AIK and KJF refined model calibrations; RJF, KS, VD, and AIK developed computational methods; AIK ran models; AIK, KJF, CCC, and VD analyzed output; AIK and KJF developed visualizations; and AIK, KJF, and CCC wrote paper; all co-authors provided feedback and approved the final manuscript.

