## Supplemental Tables for "Including variability in air temperature warming scenarios in a lake simulation model highlights uncertainty in predictions of cyanobacteria"

**Supplement A:** GLM-AED parameters used in the baseline calibrated model for Lake Mendota. See Hipsey et al. (2013; 2014; 2019) for detailed parameter descriptions and default values.

**Table A1**: Phytoplankton parameter (aed2_phyto_pars.nml) descriptions and values for the calibrated baseline GLM-AED model for Lake Mendota.

| Parameter | Description | Unit | Non-N-fixing cyano-bacteria | N-fixing cyano-bacteria | Chloro-phytes | Diatoms |
|---|---|---|---|---|---|---|
| p_initial | initial concentration of phytoplankton | mmol C $m^{-3}$ | 1 | 1 | 0.03 | 0.03 |
| p0 | minimum concentration of phytoplankton | mmol C $m^{-3}$ | 1 | 1 | 0.03 | 0.03 |
| w_p | sedimentation rate | m $d^{-1}$ | 0.5 | 0.5 | -0.10 | -0.40 |
| Ycc | carbon to chlorophyll ratio | mg C mg chl $a^{-1}$ | 50 | 50 | 40 | 40 |
| Pmax | phytoplankton max. growth rate at 20°C | $d^{-1}$ | 0.5 | 0.7 | 1.4 | 2.3 |
| fT_Method | temperature limitation function of growth | - | 1 | 1 | 1 | 1 |
| vT | Arrhenius temperature multiplier for growth | - | 1.09 | 1.05 | 1.08 | 1.08 |
| Tstd | standard temperature | °C | 20 | 20 | 20 | 11 |
| Topt | optimum temperature | °C | 30 | 28.5 | 28 | 13 |
| Tmax | maximum temperature | °C | 37 | 37 | 33 | 20 |
| lightModel | type of light response function | - | 1 | 1 | 1 | 1 |
| IK | half-saturation constant for light limitation of growth | microE $m^{-2}$ $s^{-1}$ | 33 | 35 | 70 | 10 |
| Ist | saturating light intensity (if light model=1) | microE $m^{-2}$ $s^{-1}$ | 250 | 250 | 170 | 150 |

| | | | | | | |
|---|---|---|---|---|---|---|
| KePHY | specific attenuation coefficient | mmol C $m^{-3}$ $m^{-1}$ | 0.0012 | 0.0012 | 0.0012 | 0.0012 |
| krp | fraction of primary production lost to exudation | - | 0.003 | 0.005 | 0.005 | 0.002 |
| kr | phytoplankton respiration rate at 20°C | - | 0.05 | 0.04 | 0.078 | 0.055 |
| vr | Arrhenius temperature multiplier for respiration | - | 1.1 | 1.08 | 1.05 | 1.12 |
| fres | fraction of metabolic loss that is true respiration | - | 0.75 | 0.75 | 0.75 | 0.75 |
| fdom | fraction of metabolic loss that is DOM | - | 0.25 | 0.25 | 0.25 | 0.25 |
| salTol | type of salinity limitation function | - | 0 | 0 | 0 | 0 |
| Bep | salinity limitation value at maximum salinity | | 2 | 2 | 2 | 2 |
| maxSP | maximum salinity | g $kg^{-1}$ | 35 | 35 | 35 | 35 |
| Sop | optimum salinity | g $kg^{-1}$ | 0.01 | 0.01 | 0.01 | 0.01 |
| simDINUptake | simulate DIN uptake | - | 1 | 1 | 1 | 1 |
| SimDONUptake | simulate DON uptake | - | 0 | 0 | 0 | 0 |
| simNFixation | simulate N fixation | - | 0 | 1 | 0 | 0 |
| simINDynamics | simulate internal N dynamics | - | 2 | 2 | 2 | 2 |
| N_o | minimum nitrogen concentration for uptake | mmol N $m^{-3}$ | 0 | 0 | 0 | 0 |
| KN | half-saturation concentration of nitrogen | mmol N $m^{-3}$ | 1 | 1 | 2.7 | 3.5 |
| INcon | constant internal nitrogen concentration, used if simINDynamics = 0 or 1 | mmol N mmol $C^{-1}$ | 0.035 | 0.035 | 0.035 | 0.035 |
| INmin | minimum internal nitrogen concentration, used if simINDynamics = 2 | mmol N mmol $C^{-1}$ | 0.06 | 0.06 | 0.077 | 0.077 |

| | | | | | | |
|---|---|---|---|---|---|---|
| INmax | maximum internal nitrogen concentration, used if simINDynamics = 2 | mmol N mmol $C^{-1}$ | 0.206 | 0.137 | 0.12 | 0.129 |
| UNmax | maximum nitrogen uptake, used is simINDynamics = 2 | mmol N $m^{-3}$ $d^{-1}$ | 0.068 | 0.1 | 0.051 | 0.13 |
| gthRedNFix | growth rate reduction under maximum nitrogen fixation, used if simNFixation>0 | $d^{-1}$ | 1 | 0.67 | 1 | 1 |
| NFixationRate | nitrogen fixation rate | mmol N mmol $C^{-1}$ $d^{-1}$ | 0 | 0.13 | 0 | 0 |
| simDIPUptake | simulate DIP uptake | - | 1 | 1 | 1 | 1 |
| simIPDynamics | simulate internal phosphorus dynamics | - | 2 | 2 | 2 | 2 |
| P_o | minimum phosphorus concentration for uptake | mmol P $m^{-3}$ | 0 | 0 | 0 | 0 |
| KP | half-saturation concentration of phosphorus | mmol P $m^{-3}$ | 0.05 | 0.05 | 0.07 | 0.15 |
| IPcon | constant internal phosphorus concentration, used if simIPDynamics = 0 or 1 | mmol P mmol $C^{-1}$ | 0.0015 | 0.0015 | 0.0015 | 0.0015 |
| IPmin | minimum internal phosphorus concentration, used if simIPDynamics = 2 | mmol P mmol $C^{-1}$ | 0.00077 | 0.0019 | 0.0023 | 0.0081 |
| IPmax | maximum internal phosphorus concentration, used if simIPDynamics = 2 | mmol P mmol $C^{-1}$ | 0.0089 | 0.0089 | 0.023 | 0.033 |
| UPmax | maximum phosphorus uptake, used if simIPDynamics = 2 | mmol P $m^{-3}$ $d^{-1}$ | 0.0039 | 0.0039 | 0.0027 | 0.007 |
| simSiUptake | simulate Si uptake | - | 0 | 0 | 0 | 1 |
| Si_o | minimum silica concentration for uptake | mmol Si $m^{-3}$ | 0 | 0 | 0 | 0 |

| | | | | | | |
|---|---|---|---|---|---|---|
| KSi | half-saturation concentration of silica | mmol Si $m^{-3}$ | 0 | 0 | 0 | 2.5 |
| Sicon | constant internal silica concentration | mmol Si mmol $C^{-1}$ | 0 | 0 | 0 | 0.4 |

**Table A2**: Lake physics (glm2.nml) parameter descriptions and values for the calibrated baseline GLM-AED model for Lake Mendota.

| Parameter Name | Description | Unit | Baseline value |
|---|---|---|---|
| ***glm_setup*** | | | |
| max_layers | maximum number of layers | - | 500 |
| min_layer_vol | minimum layer volume | $m^3$ | 0.025 |
| mix_layer_thick | minimum layer thickness | m | 0.125 |
| max_layer_thick | maximum layer thickness | m | 0.5 |
| Kw | background light attenuation | $m^{-1}$ | 0.15 |
| coef_mix_conv | mixing coefficient: convective overturn | - | 0.125 |
| coef_wind_stir | mixing coefficient: wind stirring | - | 0.23 |
| coef_mix_shear | mixing coefficient: shear production | - | 0.2 |
| coef_mix_turb | mixing coefficient: unsteady turbulence effects | - | 0.51 |
| coef_mix_KH | mixing coefficient: hypolimnetic Kelvin-Helmholtz turbulent billows | - | 0.3 |
| coef_mix_hyp | mixing coefficient: hypolimnetic turbulence | - | 0.5 |
| deep_mixing | flag to disable deep-mixing | - | .true. |
| ***wq_setup*** | | | |
| wq_lib | selects WQ library: 'aed2' or 'fabm' | - | 'aed2' |
| wq_nml_file | name of .nml file ot be passed to WQ library | - | 'aed2.nml' |
| ode_method | ODE numerical scheme for source and sink dynamics (see Hipsey et al. 2014 for details) | - | 1 |
| split_factor | number of biogeochemical time steps per physical time step | - | 1 |
| bioshade_feedback | feedback of bio-turbidity to temperature equation | - | .true. |
| repair_state | option to repair state variables that have -ve's | - | .true. |
| multi_ben | option for benthic fluxes on flanks of all layers | - | .true. |
| ***morphometry*** | | | |
| lake_name | name of the lake | - | 'Mendota' |
| latitude | latitude of lake | °N | 43.10 |
| longitude | longitude of lake | °E | -89.41 |
| bsn_len | basin length at crest | m | 9500 |
| bsn_wid | basin width at crest | m | 7400 |
| bsn_vals | number of depth points on height-area relationship | - | 30 |

| | | | |
|---|---|---|---|
| H | elevations, comma separated list (length=bsn_vals) | m | See Table B3 |
| A | area, comma separated list (length=bsn_vals) | $m^2$ | See Table B3 |
| ***time*** | | | |
| timefmt | method to specify start and duration of model run | - | 2 |
| start | nominal start date, format = 'yyyy-mm-dd hh:mm:ss' | - | '2003-11-08 00:00:00' |
| stop | nominal end date, format = 'yyyy-mm-dd hh:mm:ss' | - | '2014-12-31 23:00:00' |
| dt | time step for integration | s | 3600 |
| timezone | time zone | - | -6 |
| ***output*** | | | |
| out_dir | path to output directory | - | '.' |
| out_fn | name of output netcdf file | - | 'output' |
| nsave | save results every 'nsave' timesteps | - | 24 |
| ***init_profiles*** | | | |
| num_depths | number of depths provided for initial profiles | - | 6 |
| the_depths | the depths of the initial profile points | m | 0,4,8,12,16,20 |
| the_temps | the temperature of the initial profile points | °C | 12.8, 12.7, 12.7, 12.7, 12.6, 12.6 |
| the_sals | the salinity of the initial profile points | psu | 0, 0, 0, 0, 0, 0 |
| lake_depth | initial lake depth | m | 25 |
| num_wq_vars | number of non GLM (i.e. FABM) vars to be initialized | - | 13 |
| wq_names | names of non GLM (i.e. FABM) vars to be initialized | - | See Table B4 |
| wq_init_vals | array of FABM vars (rows = vars; cols = depths) | - | See Table B4 |
| ***meteorology*** | | | |
| met_sw | switch to include surface meteorological forcing | - | .true. |
| lw_type | type of longwave data supplied | - | 'LW_IN' |
| rain_sw | include rainfall nutrient composition | - | .false. |
| atm_stab | account for non-neutral atmospheric stability | - | .false. |
| catchrain | flat that enables runoff from exposed banks of lake area | - | .false. |
| rad_mode | short and long wave radiation model configuration (see Hipsey et al. 2014 for details) | - | 2 |
| albedo_mode | shortwave albedo calculation method | - | 1 |
| cloud_mode | atmospheric emissivity calculation method | - | 4 |
| meteo_fl | name of file with meteorology input data | - | 'met_hourly.csv' |

| | | | |
|---|---|---|---|
| subdaily | | - | .true. |
| wind_factor | wind multiplication factor | - | 1 |
| sw_factor | shortwave radiation multiplication factor | - | 0.9 |
| cd | bulk aerodynamic coefficient for transfer for momentum | - | 0.00106 |
| ce | bulk aerodynamic coefficient for transfer for latent heat transfer | - | 0.0014 |
| ch | bulk aerodynamic coefficient for transfer for sensible heat transfer | - | 0.0013 |
| rain_threshold | rainfall amount required before runoff from exposed banks | m | 0.01 |
| runoff_coef | conversion of rainfall to runoff in exposed banks | - | 0.3 |
| ***inflow*** | | | |
| num_inflows | number of inflowing streams | - | 3 |
| names_of_strms | names of streams, comma separated list | - | 'Highway', 'Pheasant', 'Balance' |
| strm_hf_angle | stream half angle | ° | 65,65,65 |
| strmbd_slope | streambed slope | ° | 3,3,3 |
| strmbd_drag | streambed drag coefficient | - | 0.016,0.016,0.016 |
| inflow_factor | inflow flow rate multiplier | - | 1, 1, 1 |
| inflow_fl | inflow data filenames, comma separated list | - | 'inflow_YaharaHighway.csv', 'inflow_Pheasant.csv', 'inflow_BalanceAdj.csv' |
| inflow_varnum | number of columns (excluding date) to be read | - | 13 |
| inflow_vars | variable names of inflow file columns (must include FLOW, SALT, and TEMP) | - | 'FLOW', 'SALT', 'TEMP', 'OGM_doc', 'OGM_poc','OGM_don', 'NIT_nit', 'NIT_amm', 'OGM_pon', 'PHS_frp', 'OGM_dop', 'OGM_pop', 'PHS_frp_ads' |
| ***outflows*** | | | |
| num_outlet | number of outlets | - | 1 |
| flt_off_sw | floating offtake switches | - | .false. |
| outl_elvs | outlet elevations, comma separated list | - | 256.8 |

| | | | |
|---|---|---|---|
| bsn_len_outl | basin length at outlet(s) | m | 799 |
| bsn_wid_outl | basin width at outlet(s) | m | 398 |
| outflow_fl | outflow data file | - | 'outflow.csv' |
| outflow_factor | outflow flow rate multiplier | - | 1 |
| ***snowice*** | | | |
| snow_albedo_factor | scaling factor for snow albedo | | 1 |
| snow_rho_max | maximum snow density | kg $m^{-3}$ | 500 |
| snow_rho_min | minimum snow density | kg $m^{-3}$ | 100 |
| ***sed_heat*** | | | |
| sed_temp_mean | annual mean temperature of lake sediments | °C | 9.326911 |
| sed_temp_amplitude | seasonal amplitude of sediment temperature variation | °C | 6 |
| sed_temp_peak_doy | day of year when sediment temperature peaks | d | 231 |

**Table A3**: Elevation (H) and area (A) for glm2.nml for Lake Mendota.

| **Mendota** | |
|---|---|
| **H (m)** | **A ($m^2$)** |
| 234.5 | 0 |
| 235.4 | 216000 |
| 236.2 | 821000 |
| 237.1 | 2560000 |
| 238.0 | 4040000 |
| 238.9 | 5957000 |
| 239.7 | 7777000 |
| 240.6 | 9963000 |
| 241.5 | 12271000 |
| 242.4 | 14100000 |
| 243.2 | 15659000 |
| 244.1 | 17241000 |
| 245.0 | 18990000 |
| 245.8 | 20082000 |
| 246.7 | 21564000 |
| 247.6 | 22809000 |
| 248.5 | 23789000 |
| 249.3 | 24686000 |
| 250.2 | 25311000 |
| 251.1 | 26084000 |
| 251.9 | 26745000 |
| 252.8 | 27502000 |
| 253.7 | 28064000 |
| 254.6 | 29022000 |
| 255.4 | 30154000 |
| 256.3 | 31530000 |
| 257.2 | 33404000 |
| 258.1 | 35179000 |
| 258.9 | 38308000 |
| 259.8 | 39866000 |

**Table A4:** Depth profile of Lake Mendota water quality variable initial conditions.

| | **Depth (m)** | | | | | |
|---|---|---|---|---|---|---|
| **wq_names** | **0** | **4** | **8** | **12** | **16** | **20** |
| CAR_pH | 8.3 | 8.3 | 8.3 | 8.3 | 8.1 | 8.1 |
| CAR_dic | 3746 | 3632 | 3635 | 3635 | 3781 | 3781 |
| OGM_don | 58 | 59 | 59 | 59 | 60 | 60 |
| OGM_pon | 6 | 6 | 6 | 6 | 6 | 6 |
| OGM_dop | 1.53 | 1.54 | 1.34 | 1.06 | 1.86 | 2.66 |
| OGM_pop | 0.55 | 0.55 | 0.48 | 0.38 | 0.67 | 0.96 |
| OGM_doc | 605 | 524 | 551 | 551 | 518 | 518 |
| OGM_poc | 3 | 3 | 2 | 2 | 3 | 3 |
| SIL_rsi | 28.38 | 28.66 | 22.56 | 5.642 | 41.71 | 41.71 |
| OXY_oxy | 275.0 | 275.0 | 271.9 | 268.8 | 262.5 | 259.4 |
| NIT_amm | 15.4 | 15.4 | 16.5 | 15.6 | 15.6 | 30.8 |
| NIT_nit | 4.8 | 4.8 | 8.0 | 9.1 | 9.1 | 4.8 |
| PHS_frp_ads | - | - | - | - | - | - |
| PHS_frp | 3.04 | 3.04 | 2.61 | 1.20 | 1.20 | 5.94 |

**Table A5**: Descriptions and values for water chemistry (aed2.nml) parameters in calibrated baseline GLM-AED Lake Mendota model.

| Parameter Name | Description | Unit | Baseline value |
|---|---|---|---|
| ***sed_flux*** | | | |
| sedflux_model | sediment flux model type | - | 'Constant' |
| ***sed_constant*** | | | |
| Fsed_oxy | sedimentation flux for oxygen | mmol $m^{-2}$ $d^{-1}$ | -12.55 |
| Fsed_rsi | sedimentation flux for silica | mmol $m^{-2}$ $d^{-1}$ | 0.9745 |
| Fsed_amm | sedimentation flux for ammonia | mmol $m^{-2}$ $d^{-1}$ | 22.13 |
| Fsed_nit | sedimentation flux for nitrogen | mmol $m^{-2}$ $d^{-1}$ | -8.57 |
| Fsed_frp | sedimentation flux for phosphorus | mmol $m^{-2}$ $d^{-1}$ | 0.40 |
| Fsed_don | sedimentation flux for dissolved organic nitrogen | mmol $m^{-2}$ $d^{-1}$ | 0.0 |
| Fsed_dop | sedimentation flux for dissolved organic phosphorus | mmol $m^{-2}$ $d^{-1}$ | 0.0 |
| Fsed_poc | sedimentation flux for particulate organic carbon | mmol $m^{-2}$ $d^{-1}$ | 0.0 |
| Fsed_doc | sedimentation flux for dissolved organic carbon | mmol $m^{-2}$ $d^{-1}$ | 0.0 |
| Fsed_dic | sedimentation flux for dissolved inorganic carbon | mmol $m^{-2}$ $d^{-1}$ | 3 |
| Fsed_ch4 | sedimentation flux for methane | mmol $m^{-2}$ $d^{-1}$ | 0.5 |
| Fsed_feii | sedimentation flux for iron | mmol $m^{-2}$ $d^{-1}$ | 0 |
| ***tracer*** | | | |
| num_tracers | number of tracers to model | - | 1 |
| decay | vector of decay rates for each simulated tracer group | $d^{-1}$ | 0 |
| settling | settling sub-model | - | -0.1 |
| fsed | vector of sediment flux rates for each simulated tracer group | g $(m^2*s)^{-1}$ | 0 |
| epsilon | vector of resuspension rate coefficient | g $m^{-2}$ $s^{-1}$ | 0.02 |
| tau_0 | vector of critical shear stress for resuspension | N $m^{-2}$ | 0.01 |
| tau_r | reference shear stress | N $m^{-2}$ | 1 |
| Ke_ss | vector of specific light attenuation constants for each simulated tracer group | $m^{-1}$ $(g\ m^{-3})^{-1}$ | 0.02 |
| retention_time | activates the retention time variable | - | .true. |
| ***oxygen*** | | | |
| Fsed_oxy | sediment oxygen demand | mmol $m^{-2}$ $d^{-1}$ | -60.375 |

| | | | |
|---|---|---|---|
| Ksed_oxy | half-saturation concentration of oxygen sediment flux | mmol $m^{-3}$ | 15.875 |
| theta_sed_oxy | Arrhenius temperature multiplier for sediment oxygen flux | - | 1.07 |
| oxy_min | minimum DO concentration | mmol $m^{-3}$ | 0 |
| oxy_max | maximum DO concentration | mmol $m^{-3}$ | 500 |
| ***carbon*** | | | |
| dic_initial | initial DIC concentration | mmol $m^{-3}$ | !3750 |
| Fsed_dic | sediment $CO_2$ flux | mmol $m^{-2}$ $d^{-1}$ | 4.908 |
| Ksed_dic | half-saturation oxygen concentration controlling $CO_2$ flux | mmol $m^{-3}$ | 24.34 |
| theta_sed_dic | Arrhenius temperature multiplier for sediment $CO_2$ flux | - | 1.02 |
| pH_initial | initial water column pH | - | 8.4 |
| atmco2 | atmospheric $CO_2$ concentration | ppm | 400e-6 |
| ionic | average ionic strength of the water column | meq | 0.3 |
| ch4_initial | initial $CH_4$ concentration | mmol $m^{-3}$ | 27 |
| Rch4ox | maximum reaction rate of CH4 oxidation at 20ºC | - | 0.01 |
| Kch4ox | half-saturation oxygen concentration for $CH_4$ oxidation | pp | 0.5 |
| vTch4ox | Arrhenius temperature multiplier for $CH_4$ oxidation | meq | 1.08 |
| Fsed_ch4 | sediment $CH_4$ flux | mmol $m^{-2}$ $d^{-1}$ | 10 |
| Ksed_ch4 | half-saturation oxygen concentration controlling $CH_4$ flux | mmol $m^{-3}$ | 100 |
| theta_sed_ch4 | Arrhenius temperature multiplier for sediment $CH_4$ flux | - | 1.08 |
| methane_reactant_variable | state variable to be consumed during $CH_4$ oxidation | - | 'OXY_oxy' |
| ***silica*** | | | |
| rsi_initial | | | 300 |
| Fsed_rsi | sediment flux for silica | mmol $m^{-2}$ $d^{-1}$ | 0.6 |
| Ksed_rsi | release rate for silica | mmol $m^{-3}$ | 153.5 |
| theta_sed_rsi | Arrhenius temperature multiplier for silica | - | 1.0302 |
| silica_reactant_ | link for silica reactant variable | - | 'OXY_oxy' |

| variable | | | |
|---|---|---|---|
| ***nitrogen*** | | | |
| Rnitrif | maximum nitrification rate at 20ºC | $d^{-1}$ | 0.106 |
| Rdenit | maximum denitrification rate at 20ºC | $d^{-1}$ | 0.04 |
| Fsed_amm | sediment ammonium flux | mmol $m^{-2}$ $d^{-1}$ | 22.13 |
| Fsed_nit | sediment nitrate flux | mmol $m^{-2}$ $d^{-1}$ | -9.466 |
| Knitrif | half-saturation oxygen concentration for nitrification | mmol $m^{-3}$ | 46.875 |
| Kdenit | half-saturation oxygen concentration for denitrification | mmol $m^{-3}$ | 15.24 |
| Ksed_amm | half-saturation oxygen concentration controlling NH4 flux | mmol $m^{-3}$ | 22.158 |
| Ksed_nit | half-saturation oxygen concentration controlling NO3 flux | mmol $m^{-3}$ | 15.62 |
| theta_nitrif | Arrhenius temperature multiplier for nitrification | - | 1.08 |
| theta_denit | Arrhenius temperature multiplier for denitrification | - | 1.05 |
| theta_sed_amm | Arrhenius temperature multiplier for sediment NH4 flux | - | 1.139 |
| theta_sed_nit | Arrhenius temperature multiplier for sediment NO3 flux | - | 1.055 |
| nitrif_reactant_variable | state variable to be consumed during nitrification | - | 'OXY_oxy' |
| denit_product_variable | state variable to be incremented from denitrification | - | '' |
| ***phosphorus*** | | | |
| Fsed_frp | sediment PO4 flux | mmol $m^{-2}$ $d^{-1}$ | 0.1 |
| Ksed_frp | half-saturation oxygen concentration controlling PO4 flux | mmol $m^{-3}$ | 15.6 |
| theta_sed_frp | Arrhenius temperature multiplier for sediment PO4 flux | - | 1.0324 |
| phosphorus_reactant_variable | state variable linked to sediment release | - | 'OXY_oxy' |
| simPO4Adsorption | switch to enable PO4 adsorption/desorption model | - | .true. |
| ads_use_external_tss | switch to set external environment variable as substrate | - | .false. |
| po4sorption_target_variable | variable PO4 will adsorb onto | - | '' |
| PO4AdsorptionModel | sorption algorithm to use | - | 1 |

| | | | |
|---|---|---|---|
| kpo4p | sorption constant | - | 0.1 |
| ads_use_pH | switch to engage pH dependency in sorption algorithm | - | .false. |
| Kadsratio | sorption constant | - | 0.7 |
| Qmax | sorption constant | - | 0.00016 |
| w_po4ads | settling rate of adsorbed PO4 | m day-1 | -1.2 |
| ***organic_matter*** | | | |
| pon_initial | | | 50 |
| don_initial | | | 50 |
| w_pon | settling rate for particulate organic nitrogen | m day-1 | -0.1238 |
| Rpon_miner | hydrolysis/breakdown rate of particulate organic N pool at 20ºC | $d^{-1}$ | 0.035 |
| Rdon_miner | hydrolysis/breakdown rate of dissolved organic N pool at 20ºC | $d^{-1}$ | 0.01 |
| Fsed_pon | sediment flux of particulate organic nitrogen | mmol $m^{-2}$ $d^{-1}$ | 0 |
| Fsed_don | sediment flux of dissolved organic nitrogen | mmol $m^{-2}$ $d^{-1}$ | 0.1 |
| Kpon_miner | half-saturation oxygen concentration controlling PON hydrolysis | mmol $m^{-3}$ | 62.5 |
| Kdon_miner | half-saturation oxygen concentration controlling DON hydrolysis | mmol $m^{-3}$ | 62.5 |
| Ksed_don | half-saturation oxygen concentration controlling DON sediment flux | mmol $m^{-3}$ | 4.5 |
| theta_pon_miner | Arrhenius temperature multiplier for PON breakdown | - | 1.08 |
| theta_don_miner | Arrhenius temperature multiplier for DON breakdown | - | 1.08 |
| theta_sed_don | Arrhenius temperature multiplier for sediment flux of DON | - | 1.08 |
| don_miner_product_variable | state variable to be the product of DON mineralization | - | 'NIT_amm' |
| pop_initial | initial particulate organic phosphorus concentration | mmol $m^{-3}$ | 0.5 |
| dop_initial | initial dissolved organic phosphorus concentration | mmol $m^{-3}$ | 0.5 |
| w_pop | settling rate of particulate organic phosphorus | m $d^{-1}$ | -1 |

| | | | |
|---|---|---|---|
| Rpop_miner | hydrolysis/breakdown rate of particulate organic P pool at 20ºC | $d^{-1}$ | 0.03 |
| Rdop_miner | hydrolysis/breakdown rate of dissolved organic P pool at 20ºC | $d^{-1}$ | 0.01 |
| Fsed_pop | sediment flux of particulate organic phosphorus | mmol $m^{-2}$ $d^{-1}$ | -0.01 |
| Fsed_dop | sediment flux of dissolved organic phosphorus | mmol $m^{-2}$ $d^{-1}$ | 0.03 |
| Kpop_miner | half-saturation oxygen concentration controlling POP hydrolysis | mmol $m^{-3}$ | 62.5 |
| Kdop_miner | half-saturation oxygen concentration controlling DOP hydrolysis | mmol $m^{-3}$ | 62.5 |
| Ksed_dop | half-saturation oxygen concentration controlling DOP sediment flux | mmol $m^{-3}$ | 40.5 |
| theta_pop_miner | Arrhenius temperature multiplier for POP breakdown | - | 1.08 |
| theta_dop_miner | Arrhenius temperature multiplier for DOP breakdown | - | 1.08 |
| theta_sed_dop | Arrhenius temperature multiplier for sediment flux of DOP | - | 1.08 |
| dop_miner_product_variable | state variable to be the product of DOP mineralization | - | 'PHS_frp' |
| poc_initial | initial particulate organic carbon concentration | mmol $m^{-3}$ | 80 |
| doc_initial | initial dissolved organic carbon concentration | mmol $m^{-3}$ | 400 |
| w_poc | settling rate of particulate organic carbon | m day-1 | -0.1238 |
| Rpoc_miner | hydrolysis/breakdown rate of particulate organic C pool at 20ºC | $d^{-1}$ | 0.04 |
| Rdoc_miner | hydrolysis/breakdown rate of dissolved organic C pool at 20ºC | $d^{-1}$ | 0.003 |
| Fsed_poc | sediment flux of particulate organic carbon | mmol $m^{-2}$ $d^{-1}$ | -0.01 |
| Fsed_doc | sediment flux of dissolved organic carbon | mmol $m^{-2}$ $d^{-1}$ | 0.044 |
| Kpoc_miner | half-saturation oxygen concentration controlling POC hydrolysis | mmol $m^{-3}$ | 62.5 |
| Kdoc_miner | half-saturation oxygen concentration controlling DOC hydrolysis | mmol $m^{-3}$ | 62.5 |

| | | | |
|---|---|---|---|
| Ksed_doc | half-saturation oxygen concentration controlling DOC sediment flux | mmol $m^{-3}$ | 16.81 |
| theta_poc_miner | Arrhenius temperature multiplier for POC breakdown | - | 1.08 |
| theta_doc_miner | Arrhenius temperature multiplier for DOC breakdown | - | 1.08 |
| theta_sed_doc | Arrhenius temperature multiplier for sediment flux of DOC | - | 1.08 |
| KeDOM | specific light attenuation for dissolved organic matter | $m^{-1}$ | 0.001 |
| KePOM | specific light attenuation for particulate organic matter | $m^{-1}$ | 0.001 |
| doc_miner_reactant_variable | state variable to be linked to rate of DOC mineralization | - | 'OXY_oxy' |
| doc_miner_product_variable | state variable to be the product of DOC mineralization | - | '' |
| ***phytoplankton*** | | | |
| num_phytos | number of phytoplankton groups within this module to include | - | 4 |
| the_phytos | list of ID's of groups in aed_phyto_pars.nml | - | 1,2,3,4 |
| p_excretion_target_variable | state variable to receive P from excretion | - | 'OGM_dop' |
| n_excretion_target_variable | state variable to receive N from excretion | - | 'OGM_don' |
| c_excretion_target_variable | state variable to receive C from excretion | - | 'OGM_doc' |
| si_excretion_target_variable | state variable to receive Si from excretion | - | ' ' |
| p_mortality_target_variable | state variable to receive P from mortality | - | 'OGM_pop' |
| n_mortality_target_variable | state variable to receive N from mortality | - | 'OGM_pon' |
| c_mortality_target_variable | state variable to receive C from mortality | - | 'OGM_poc' |
| si_mortality_target_variable | state variable to receive Si from mortality | - | ' ' |
| p1_uptake_target_variable | state variable to be linked for P uptake | - | 'PHS_frp' |
| n1_uptake_target_variable | state variable to be linked for N1 uptake | - | 'NIT_nit' |
| n2_uptake_target_variable | state variable to be linked for N2 uptake | - | 'NIT_amm' |
| si_uptake_target_variable | state variable to be linked for Si uptake | - | 'SIL_rsi' |
| do_uptake_target_variable | state variable to be linked for DO uptake | - | 'OXY_oxy' |

| | | | |
|---|---|---|---|
| c_uptake_target_variable | state variable to be linked for C uptake | - | ' ' |
| ***zooplankton*** | | | |
| num_zoops | number of zooplankton groups within this module to include | - | 3 |
| the_zoops | list of ID's of groups in aed_zoop_pars.nml | - | 1,2,3 |
| dn_target_variable | state variable linked to provide/receive dissolved organic nitrogen | - | 'OGM_don' |
| pn_target_variable | state variable linked to provide/receive particulate organic nitrogen | - | 'OGM_pon' |
| dp_target_variable | state variable linked to provide/receive dissolved organic phosphorus | - | 'OGM_dop' |
| pp_target_variable | state variable linked to provide/receive particulate organic phosphorus | - | 'OGM_pop' |
| dc_target_variable | state variable linked to provide/receive dissolved organic carbon | - | 'OGM_doc' |
| pc_target_variable | state variable linked to provide/receive particulate organic carbon | - | 'OGM_poc' |
| ***totals*** | | | |
| TN_vars | all variables that are included in TN | - | NIT_nit |
| | | - | NIT_amm |
| | | - | OGM_don |
| | | - | OGM_pon |
| | | - | PHY_CYANOPCH1_IN |
| | | - | PHY_CYANONPCH2_IN |
| | | - | PHY_CHLOROPCH3_IN |
| | | - | PHY_DIATOMPCH4_IN |
| TN_varscale | scaling of TN variables' contribution | - | 1, 1, 1, 1, 0.15 |
| TP_vars | all variables that are included in TP | - | PHS_frp |
| | | - | PHS_frp_ads |

| | | | |
|---|---|---|---|
| | | - | OGM_dop |
| | | - | OGM_pop |
| | | - | PHY_CYANOPCH1 |
| | | - | PHY_CYANONPCH2 |
| | | - | PHY_CHLOROPCH3 |
| | | - | PHY_DIATOMPCH4 |
| TP_varscale | scaling of TP variables' contribution | - | 1, 1, 0.7, 0.7, 0.01, 0.01, 0.01, 0.01 |
| TOC_vars | all variables that are included in TOC | - | OGM_doc |
| | | - | OGM_poc |
| TOC_varscale | scaling of TOC variables' contribution | - | 1, 1, 1, 1, 1, 1 |
| TSS_vars | all variables that are included in TSS | - | TRC_ssl |

**Table A6**: Zooplankton parameter (aed2_zoop_pars.nml) descriptions and values for calibrated baseline models for Lake Mendota, derived from Kara et al. (2012).

| Parameter | Description | Unit | Copepods | Large cladocerans | Small cladocerans |
|---|---|---|---|---|---|
| zoop_initial | initial concentration of zooplankton | mmol $m^{-3}$ | 5 | 5 | 5 |
| min_zoo | minimum concentration of zooplankton | mmol $m^{-3}$ | 0.1 | 0.1 | 0.1 |
| Rgrz_zoo | zooplankton grazing rate | $d^{-1}$ | 0.67 | 0.53 | 0.22 |
| fassim_zoo | assimilation efficiency for zooplankton grazing | - | 0.8 | 0.7 | 0.8 |
| K_grz_zoo | half-saturation constant for zooplankton grazing | mmol $m^{-3}$ | 167 | 167 | 167 |
| theta_grz_zoo | Arrhenius temperature multiplier for zooplankton grazing | - | 1.08 | 1.08 | 1.08 |
| Rresp_zoo | respiration rate coefficient | $d^{-1}$ | 0.1 | 0.2 | 0.075 |
| Rmort_zoo | mortality rate coefficient | $d^{-1}$ | 0.018 | 0.034 | 0.012 |
| ffecal_zoo | fecal pellet fraction of respiration | - | 0.2 | 0.2 | 0.2 |
| fexcr_zoo | excretion fraction of respiration | - | 0.7 | 0.7 | 0.7 |
| ffecal_sed | fraction of fecal pellets that sink directly to sediments | - | 0.7 | 0.1 | 0.1 |
| theta_resp_zoo | Arrhenius temperature multiplier for zooplankton respiration | - | 1.07 | 1.07 | 1.07 |
| Tstd_zoo | standard temperature for zooplankton respiration temperature dependence | °C | 20 | 20 | 20 |
| Topt_zoo | optimum temperature for zooplankton respiration temperature dependence | °C | 19 | 20 | 18 |

**Supplement B:** Results of Anderson-Darling statistical tests.

**Table B1.** Results of Anderson-Darling tests comparing across-replicate maximum daily water temperatures between the baseline scenario (+0°C) and each of the warming distribution scenarios across air temperature offsets (+1°C to +5°C). The Anderson-Darling criterion (AD), standardized test statistic (T.AD), and *P*-values (asymptotic approximation) are reported for each pairwise scenario comparison. *P*-values in bold indicate significant (Bonferroni-corrected α = 0.01) shifts in the distribution; values in italics indicate shifts with marginal significance (Bonferroni-corrected α = 0.02).

| **Temp. Offset** | **Uniform** | | | **Normal** | | | **Poisson- Tail 2** | | | **Poisson- Tail 4** | | | **Poisson- Tail 8** | | |
|---|---|---|---|---|---|---|---|---|---|---|---|---|---|---|---|
| | **AD** | **T.AD** | ***P*** | **AD** | **T. AD** | ***P*** | **AD** | **T. AD** | ***P*** | **AD** | **T. AD** | ***P*** | **AD** | **T. AD** | ***P*** |
| +1°C | 13.33 | 16.20 | **0.00** | 33.82 | 43.13 | **0.00** | 30.73 | 39.08 | **0.00** | 28.57 | 36.23 | **0.00** | 27.28 | 34.54 | **0.00** |
| +2°C | 50.73 | 65.37 | **0.00** | 84.27 | 109.45 | **0.00** | 98.00 | 127.49 | **0.00** | 85.54 | 111.11 | **0.00** | 79.89 | 103.69 | **0.00** |
| +3°C | 111.38 | 145.07 | **0.00** | 156.76 | 204.72 | **0.00** | 186.33 | 243.58 | **0.00** | 167.55 | 218.90 | **0.00** | 157.64 | 205.88 | **0.00** |
| +4°C | 187.99 | 245.76 | **0.00** | 238.50 | 312.15 | **0.00** | 288.15 | 377.41 | **0.00** | 269.69 | 353.14 | **0.00** | 253.67 | 332.09 | **0.00** |
| +5°C | 274.71 | 359.74 | **0.00** | 331.89 | 434.90 | **0.00** | 403.59 | 529.13 | **0.00** | 372.52 | 488.30 | **0.00** | 355.59 | 466.05 | **0.00** |

**Table B2.** Results of Anderson-Darling tests comparing across-replicate median daily water temperatures between the baseline scenario (+0°C) and each of the warming distribution scenarios across air temperature offsets (+1°C to +5°C). The Anderson-Darling criterion (AD), standardized test statistic (T.AD), and *P*-values (asymptotic approximation) are reported for each pairwise scenario comparison. *P*-values in bold indicate significant (Bonferroni-corrected α = 0.01) shifts in the distribution; values in italics indicate shifts with marginal significance (Bonferroni-corrected α = 0.02).

| Temp. Offset | Uniform | | | Normal | | | Poisson- Tail 2 | | | Poisson- Tail 4 | | | Poisson- Tail 8 | | |
|---|---|---|---|---|---|---|---|---|---|---|---|---|---|---|---|
| | AD | T.AD | *P* | AD | T. AD | *P* | AD | T. AD | *P* | AD | T. AD | *P* | AD | T. AD | *P* |
| +1°C | 13.33 | 16.20 | **0.00** | 12.96 | 15.72 | **0.00** | 13.57 | 16.52 | **0.00** | 13.04 | 15.82 | **0.00** | 13.42 | 16.32 | **0.00** |
| +2°C | 50.73 | 65.37 | **0.00** | 51.52 | 66.40 | **0.00** | 51.65 | 66.57 | **0.00** | 50.72 | 65.34 | **0.00** | 52.03 | 67.06 | **0.00** |
| +3°C | 111.4 | 145.1 | **0.00** | 112.5 | 146.6 | **0.00** | 107.8 | 140.4 | **0.00** | 110.7 | 144.2 | **0.00** | 112.1 | 146.0 | **0.00** |
| +4°C | 188.0 | 245.8 | **0.00** | 186.4 | 243.6 | **0.00** | 179.9 | 235.1 | **0.00** | 187.5 | 245.1 | **0.00** | 186.6 | 244.0 | **0.00** |
| +5°C | 274.7 | 359.7 | **0.00** | 277.4 | 363.3 | **0.00** | 264.3 | 346.1 | **0.00** | 271.2 | 355.1 | **0.00** | 274.2 | 359.0 | **0.00** |

**Table B3.** Results of Anderson-Darling tests comparing across-replicate median daily water temperatures between the uniform distribution and each of the variable distributions (Normal, Poisson- 2, 4, or 8 Tail, denoted by Distrib.) for each non-zero air temperature offset. The Anderson-Darling criterion (AD), standardized test statistic (T.AD), and *P*-values (asymptotic approximation) are reported for each pairwise scenario comparison. *P*-values in bold indicate significant (Bonferroni-corrected α = 0.01) shifts in the distribution; values in italics indicate shifts with marginal significance (Bonferroni-corrected α = 0.02).

| **Distrib.** | | **+0** | | | **+1** | | | **+2** | | | **+3** | | | **+4** | | | **+5** | |
|---|---|---|---|---|---|---|---|---|---|---|---|---|---|---|---|---|---|---|
| | **AD** | **T.AD** | ***P*** | **AD** | **T.AD** | ***P*** | **AD** | **T.AD** | ***P*** | **AD** | **T.AD** | ***P*** | **AD** | **T.AD** | ***P*** | **AD** | **T.AD** | ***P*** |
| Normal | 0.11 | -1.17 | 1.00 | 0.14 | -1.14 | 1.00 | 0.08 | -1.21 | 1.00 | 0.08 | -1.21 | 1.00 | 0.07 | -1.22 | 1.00 | 0.06 | -1.24 | 1.00 |
| Poisson-Tail 2 | 0.00 | -1.31 | 1.00 | 0.09 | -1.19 | 1.00 | 0.10 | -1.18 | 1.00 | 0.13 | -1.14 | 1.00 | 0.25 | -0.99 | 0.97 | 0.33 | -0.88 | 0.92 |
| Poisson-Tail 4 | 0.00 | -1.31 | 1.00 | 0.10 | -1.18 | 1.00 | 0.08 | -1.21 | 1.00 | 0.07 | -1.22 | 1.00 | 0.07 | -1.22 | 1.00 | 0.19 | -1.07 | 0.99 |
| Poisson-Tail 8 | 0.00 | -1.31 | 1.00 | 0.09 | -1.19 | 1.00 | 0.09 | -1.19 | 1.00 | 0.07 | -1.23 | 1.00 | 0.07 | -1.23 | 1.00 | 0.08 | -1.21 | 1.00 |

**Table B4.** Results of Anderson-Darling tests for across-replicate maximum daily water temperatures between the uniform distribution and each of the variable distributions (Normal, Poisson- 2, 4, or 8 Tail, denoted by Distrib.) for each non-zero air temperature offset. The Anderson-Darling criterion (AD), standardized test statistic (T.AD), and *P*-values (asymptotic approximation) are reported for each pairwise scenario comparison. *P*-values in bold indicate significant (Bonferroni-corrected α = 0.01) shifts in the distribution; values in italics indicate shifts with marginal significance (Bonferroni-corrected α = 0.02).

| **Distrib.** | | **+0** | | | **+1** | | | **+2** | | | **+3** | | | **+4** | | | **+5** | |
|---|---|---|---|---|---|---|---|---|---|---|---|---|---|---|---|---|---|---|
| | **AD** | **T.AD** | ***P*** | **AD** | **T.AD** | ***P*** | **AD** | **T.AD** | ***P*** | **AD** | **T.AD** | ***P*** | **AD** | **T.AD** | ***P*** | **AD** | **T.AD** | ***P*** |
| Normal | 4.81 | 5.01 | **0.00** | 5.08 | 5.36 | **0.00** | 5.35 | 5.72 | **0.00** | 5.28 | 5.62 | **0.00** | 5.33 | 5.69 | **0.00** | 5.36 | 5.73 | **0.00** |
| Poisson-Tail 2 | 0.00 | -1.31 | 1.00 | 3.97 | 3.91 | **0.01** | 9.64 | 11.36 | **0.00** | 14.44 | 17.66 | **0.00** | 19.78 | 24.68 | **0.00** | 29.7 | 37.7 | **0.00** |
| Poisson-Tail 4 | 0.00 | -1.31 | 1.00 | 3.16 | 2.84 | *0.02* | 5.68 | 6.14 | **0.00** | 8.39 | 9.71 | **0.00** | 13.35 | 16.23 | **0.00** | 16.5 | 20.3 | **0.00** |
| Poisson-Tail 8 | 0.00 | -1.31 | 1.00 | 2.72 | 2.26 | 0.04 | 4.16 | 4.16 | **0.01** | 5.74 | 6.23 | **0.00** | 8.75 | 10.19 | **0.00** | 10.7 | 12.8 | **0.00** |

**Table B5.** Results of Anderson-Darling tests comparing median non-N-fixing cyanobacterial bloom days per year between the baseline scenario (+0°C) and each of the warming distribution scenarios across air temperature offsets (+1°C to +5°C). The Anderson-Darling criterion (AD), standardized test statistic (T.AD), and *P*-values (asymptotic approximation) are reported for each pairwise scenario comparison. *P*-values in bold indicate significant (Bonferroni-corrected α = 0.01) shifts in the distribution; values in italics indicate shifts with marginal significance (Bonferroni-corrected α = 0.02).

| **Temp. Offset** | **Uniform** | | | **Normal** | | | **Poisson- Tail 2** | | | **Poisson- Tail 4** | | | **Poisson- Tail 8** | | |
|---|---|---|---|---|---|---|---|---|---|---|---|---|---|---|---|
| | **AD** | **T.AD** | ***P*** | **AD** | **T. AD** | ***P*** | **AD** | **T. AD** | ***P*** | **AD** | **T. AD** | ***P*** | **AD** | **T. AD** | ***P*** |
| +1°C | 3.90 | 4.17 | **0.01** | 1.85 | 1.23 | 0.10 | 2.00 | 1.43 | 0.08 | 2.88 | 2.70 | 0.03 | 3.29 | 3.29 | *0.02* |
| +2°C | 0.69 | -0.45 | 0.59 | 2.31 | 1.89 | 0.05 | 2.42 | 2.04 | 0.05 | 2.14 | 1.63 | 0.07 | 2.39 | 1.99 | 0.05 |
| +3°C | 0.29 | -1.03 | 0.99 | 0.76 | -0.35 | 0.53 | 2.10 | 1.58 | 0.07 | 1.66 | 0.95 | 0.13 | 1.50 | 0.72 | 0.17 |
| +4°C | 1.69 | 0.99 | 0.13 | 0.84 | -0.23 | 0.47 | 1.40 | 0.57 | 0.19 | 1.57 | 0.83 | 0.15 | 1.74 | 1.07 | 0.12 |
| +5°C | 0.45 | -0.79 | 0.85 | 0.42 | -0.84 | 0.89 | 0.70 | -0.43 | 0.58 | 0.61 | -0.56 | 0.67 | 0.43 | -0.82 | 0.87 |

**Table B6.** Results of Anderson-Darling tests comparing median non-N-fixing cyanobacterial bloom days per year between the uniform distribution and each of the variable distributions (Normal, Poisson- 2, 4, or 8 Tail, denoted by Distrib.) for each non-zero air temperature offset. The Anderson-Darling criterion (AD), standardized test statistic (T.AD), and *P*-values (asymptotic approximation) are reported for each pairwise scenario comparison. *P*-values in bold indicate significant (Bonferroni-corrected α = 0.01) shifts in the distribution; values in italics indicate shifts with marginal significance (Bonferroni-corrected α = 0.02).

| **Distrib.** | | **+0** | | | **+1** | | | **+2** | | | **+3** | | | **+4** | | | **+5** | |
|---|---|---|---|---|---|---|---|---|---|---|---|---|---|---|---|---|---|---|
| | **AD** | **T.AD** | ***P*** | **AD** | **T.AD** | ***P*** | **AD** | **T.AD** | ***P*** | **AD** | **T.AD** | ***P*** | **AD** | **T.AD** | ***P*** | **AD** | **T.AD** | ***P*** |
| Normal | 0.26 | -1.07 | 0.99 | 0.21 | -1.14 | 1.00 | 0.49 | -0.74 | 0.81 | 0.61 | -0.56 | 0.67 | 0.34 | -0.94 | 0.95 | 0.66 | -0.49 | 0.62 |
| Poisson-Tail 2 | 0.00 | -1.44 | 1.00 | 0.32 | -0.97 | 0.97 | 0.47 | -0.76 | 0.83 | 0.51 | -0.70 | 0.78 | 0.28 | -1.04 | 0.99 | 0.27 | -1.04 | 0.99 |
| Poisson-Tail 4 | 0.00 | -1.44 | 1.00 | 0.77 | -0.33 | 0.52 | 0.33 | -0.96 | 0.96 | 0.62 | -0.55 | 0.66 | 0.57 | -0.61 | 0.71 | 0.51 | -0.70 | 0.78 |
| Poisson-Tail 8 | 0.00 | -1.44 | 1.00 | 0.25 | -1.07 | 1.00 | 0.47 | -0.76 | 0.83 | 0.86 | -0.20 | 0.45 | 0.61 | -0.55 | 0.67 | 0.58 | -0.60 | 0.70 |

**Table B7.** Results of Anderson-Darling tests comparing median N-fixing cyanobacterial bloom days per year between the baseline scenario (+0°C) and each of the warming distribution scenarios across air temperature offsets (+1°C to +5°C). The Anderson-Darling criterion (AD), standardized test statistic (T.AD), and *P*-values (asymptotic approximation) are reported for each pairwise scenario comparison. *P*-values in bold indicate significant (Bonferroni-corrected α = 0.01) shifts in the distribution; values in italics indicate shifts with marginal significance (Bonferroni-corrected α = 0.02).

| Temp. Offset | Uniform | | | Normal | | | Poisson- Tail 2 | | | Poisson- Tail 4 | | | Poisson- Tail 8 | | |
|---|---|---|---|---|---|---|---|---|---|---|---|---|---|---|---|
| | AD | T.AD | *P* | AD | T. AD | *P* | AD | T. AD | *P* | AD | T. AD | *P* | AD | T. AD | *P* |
| +1°C | 0.39 | -0.87 | 0.91 | 0.58 | -0.61 | 0.71 | 0.28 | -1.03 | 0.99 | 0.27 | -1.04 | 0.99 | 0.23 | -1.11 | 1.00 |
| +2°C | 0.88 | -0.17 | 0.43 | 0.49 | -0.73 | 0.80 | 0.39 | -0.87 | 0.91 | 0.44 | -0.81 | 0.86 | 0.27 | -1.06 | 0.99 |
| +3°C | 1.22 | 0.32 | 0.25 | 0.99 | -0.02 | 0.37 | 0.43 | -0.82 | 0.87 | 0.73 | -0.38 | 0.55 | 0.43 | -0.83 | 0.88 |
| +4°C | 0.66 | -0.49 | 0.62 | 1.11 | 0.15 | 0.30 | 1.07 | 0.10 | 0.32 | 0.40 | -0.86 | 0.90 | 0.28 | -1.04 | 0.99 |
| +5°C | 1.11 | 0.16 | 0.30 | 0.31 | -1.00 | 0.98 | 0.87 | -0.19 | 0.44 | 0.90 | -0.15 | 0.42 | 0.62 | -0.54 | 0.66 |

**Table B8.** Results of Anderson-Darling tests comparing median N-fixing cyanobacterial bloom days per year between the uniform distribution and each of the variable distributions (Normal, Poisson- 2, 4, or 8 Tail, denoted by Distrib.) for each non-zero air temperature offset. The Anderson-Darling criterion (AD), standardized test statistic (T.AD), and *P*-values (asymptotic approximation) are reported for each pairwise scenario comparison. *P*-values in bold indicate significant (Bonferroni-corrected α = 0.01) shifts in the distribution; values in italics indicate shifts with marginal significance (Bonferroni-corrected α = 0.02).

| **Distrib.** | | **+0** | | | **+1** | | | **+2** | | | **+3** | | | **+4** | | | **+5** | |
|---|---|---|---|---|---|---|---|---|---|---|---|---|---|---|---|---|---|---|
| | **AD** | **T.AD** | ***P*** | **AD** | **T.AD** | ***P*** | **AD** | **T.AD** | ***P*** | **AD** | **T.AD** | ***P*** | **AD** | **T.AD** | ***P*** | **AD** | **T.AD** | ***P*** |
| Normal | 0.26 | -1.07 | 0.99 | 0.21 | -1.14 | 1.00 | 0.49 | -0.74 | 0.81 | 0.61 | -0.56 | 0.67 | 0.34 | -0.94 | 0.95 | 0.66 | -0.49 | 0.62 |
| Poisson-Tail 2 | 0.00 | -1.44 | 1.00 | 0.32 | -0.97 | 0.97 | 0.47 | -0.76 | 0.83 | 0.51 | -0.70 | 0.78 | 0.28 | -1.04 | 0.99 | 0.27 | -1.04 | 0.99 |
| Poisson-Tail 4 | 0.00 | -1.44 | 1.00 | 0.77 | -0.33 | 0.52 | 0.33 | -0.96 | 0.96 | 0.62 | -0.55 | 0.66 | 0.57 | -0.61 | 0.71 | 0.51 | -0.70 | 0.78 |
| Poisson-Tail 8 | 0.00 | -1.44 | 1.00 | 0.25 | -1.07 | 1.00 | 0.47 | -0.76 | 0.83 | 0.86 | -0.20 | 0.45 | 0.61 | -0.55 | 0.67 | 0.58 | -0.60 | 0.70 |

**Table B9.** Results of Anderson-Darling tests for maximum N-fixing cyanobacterial bloom days between the uniform distribution and each of the variable distributions (Normal, Poisson- 2, 4, or 8 Tail, denoted by "Distrib." D) for each air temperature offset. The Anderson-Darling criterion (AD), standardized test statistic (T.AD), and *P*-values (asymptotic approximation) are reported for each pairwise scenario comparison. *P*-values in bold indicate significant (Bonferroni-corrected α = 0.01) shifts in the distribution; values in italics indicate shifts with marginal significance (Bonferroni-corrected α = 0.02).

| **Distrib.** | | **+0** | | | **+1** | | | **+2** | | | **+3** | | | **+4** | | | **+5** | |
|---|---|---|---|---|---|---|---|---|---|---|---|---|---|---|---|---|---|---|
| | **AD** | **T.AD** | ***P*** | **AD** | **T.AD** | ***P*** | **AD** | **T.AD** | ***P*** | **AD** | **T.AD** | ***P*** | **AD** | **T.AD** | ***P*** | **AD** | **T.AD** | ***P*** |
| Normal | 3.38 | 3.42 | **0.01** | 4.52 | 5.05 | **0.00** | 1.92 | 1.32 | 0.09 | 2.92 | 2.76 | *0.02* | 1.59 | 0.84 | 0.15 | 0.94 | -0.08 | 0.39 |
| Poisson-Tail 2 | 0.00 | -1.44 | 1.00 | 5.20 | 6.03 | **0.00** | 1.53 | 0.77 | 0.16 | 3.36 | 3.38 | **0.01** | 2.07 | 1.54 | 0.07 | 0.68 | -0.45 | 0.60 |
| Poisson-Tail 4 | 0.00 | -1.44 | 1.00 | 4.01 | 4.33 | **0.01** | 1.53 | 0.76 | 0.16 | 3.80 | 4.02 | **0.01** | 0.75 | -0.36 | 0.54 | 1.13 | 0.19 | 0.29 |
| Poisson-Tail 8 | 0.00 | -1.44 | 1.00 | 4.33 | 4.78 | **0.00** | 1.29 | 0.41 | 0.23 | 3.07 | 2.98 | *0.02* | 0.59 | -0.58 | 0.69 | 1.11 | 0.16 | 0.30 |

**Table B10.** Results of Anderson-Darling tests for maximum non-N-fixing cyanobacterial bloom days between the uniform distribution and each of the variable distributions (Normal, Poisson- 2, 4, or 8 Tail, denoted by Distrib.) for each non-zero air temperature offset. The Anderson-Darling criterion (AD), standardized test statistic (T.AD), and *P*-values (asymptotic approximation) are reported for each pairwise scenario comparison. *P*-values in bold indicate significant (Bonferroni-corrected α = 0.01) shifts in the distribution; values in italics indicate shifts with marginal significance (Bonferroni-corrected α = 0.02).

| **Distrib.** | | **+0** | | | **+1** | | | **+2** | | | **+3** | | | **+4** | | | **+5** | |
|---|---|---|---|---|---|---|---|---|---|---|---|---|---|---|---|---|---|---|
| | **AD** | **T.AD** | ***P*** | **AD** | **T.AD** | ***P*** | **AD** | **T.AD** | ***P*** | **AD** | **T.AD** | ***P*** | **AD** | **T.AD** | ***P*** | **AD** | **T.AD** | ***P*** |
| Normal | 2.64 | 2.35 | 0.04 | 2.34 | 1.92 | 0.05 | 5.11 | 5.91 | **0.00** | 2.63 | 2.34 | 0.04 | 1.31 | 0.45 | 0.22 | 3.47 | 3.55 | **0.01** |
| Poisson-Tail 2 | 0.00 | -1.44 | 1.00 | 3.23 | 3.20 | *0.02* | 2.75 | 2.51 | 0.03 | 5.50 | 6.46 | **0.00** | 4.23 | 4.64 | **0.01** | 3.47 | 3.55 | **0.01** |
| Poisson-Tail 4 | 0.00 | -1.44 | 1.00 | 2.02 | 1.46 | 0.08 | 4.59 | 5.15 | **0.00** | 5.52 | 6.49 | **0.00** | 0.91 | -0.13 | 0.41 | 1.78 | 1.13 | 0.11 |
| Poisson-Tail 8 | 0.00 | -1.44 | 1.00 | 2.25 | 1.79 | 0.06 | 5.29 | 6.17 | **0.00** | 3.50 | 3.59 | **0.01** | 1.26 | 0.37 | 0.24 | 2.69 | 2.42 | 0.03 |

**Table B11.** Results of Anderson-Darling tests comparing maximum non-N-fixing cyanobacterial bloom days per year between the baseline scenario (+0°C) and each of the warming distribution scenarios across air temperature offsets (+1°C to +5°C). The Anderson-Darling criterion (AD), standardized test statistic (T.AD), and *P*-values (asymptotic approximation) are reported for each pairwise scenario comparison. *P*-values in bold indicate significant (Bonferroni-corrected α = 0.01) shifts in the distribution; values in italics indicate shifts with marginal significance (Bonferroni-corrected α = 0.02).

| Temp. Offset | Uniform | | | Normal | | | Poisson- Tail 2 | | | Poisson- Tail 4 | | | Poisson- Tail 8 | | |
|---|---|---|---|---|---|---|---|---|---|---|---|---|---|---|---|
| | AD | T.AD | *P* | AD | T. AD | *P* | AD | T. AD | *P* | AD | T. AD | *P* | AD | T. AD | *P* |
| +1°C | 3.90 | 4.17 | **0.01** | 7.66 | 9.57 | **0.00** | 8.02 | 10.09 | **0.00** | 7.34 | 9.11 | **0.00** | 7.61 | 9.49 | **0.00** |
| +2°C | 0.69 | -0.45 | 0.59 | 7.36 | 9.13 | **0.00** | 5.59 | 6.59 | **0.00** | 7.21 | 8.92 | **0.00** | 7.54 | 9.39 | **0.00** |
| +3°C | 0.29 | -1.03 | 0.99 | 3.99 | 4.29 | **0.01** | 6.45 | 7.82 | **0.00** | 6.32 | 7.64 | **0.00** | 4.55 | 5.10 | **0.00** |
| +4°C | 1.69 | 0.99 | 0.13 | 4.45 | 4.96 | **0.00** | 7.91 | 9.93 | **0.00** | 4.11 | 4.46 | **0.01** | 5.41 | 6.33 | **0.00** |
| +5°C | 0.45 | -0.79 | 0.85 | 3.74 | 3.94 | **0.01** | 3.63 | 3.78 | **0.01** | 1.73 | 1.05 | 0.12 | 2.67 | 2.40 | 0.03 |

**Table B12.** Results of Anderson-Darling tests comparing maximum N-fixing cyanobacterial bloom days per year between the baseline scenario (+0°C) and each of the warming distribution scenarios across air temperature offsets (+1°C to +5°C). The Anderson-Darling criterion (AD), standardized test statistic (T.AD), and *P*-values (asymptotic approximation) are reported for each pairwise scenario comparison. *P*-values in bold indicate significant (Bonferroni-corrected α = 0.01) shifts in the distribution; values in italics indicate shifts with marginal significance (Bonferroni-corrected α = 0.02).

| Temp. Offset | Uniform | | | Normal | | | Poisson- Tail 2 | | | Poisson- Tail 4 | | | Poisson- Tail 8 | | |
|---|---|---|---|---|---|---|---|---|---|---|---|---|---|---|---|
| | AD | T.AD | *P* | AD | T. AD | *P* | AD | T. AD | *P* | AD | T. AD | *P* | AD | T. AD | *P* |
| +1°C | 0.39 | -0.87 | 0.91 | 3.31 | 3.32 | *0.02* | 4.41 | 4.89 | **0.00** | 3.36 | 3.39 | **0.01** | 3.77 | 3.98 | **0.01** |
| +2°C | 0.88 | -0.17 | 0.43 | 3.48 | 3.56 | **0.01** | 3.21 | 3.18 | *0.02* | 3.18 | 3.14 | *0.02* | 3.50 | 3.59 | **0.01** |
| +3°C | 1.22 | 0.32 | 0.25 | 3.05 | 2.94 | *0.02* | 4.95 | 5.67 | **0.00** | 4.62 | 5.20 | **0.00** | 2.80 | 2.58 | 0.03 |
| +4°C | 0.66 | -0.49 | 0.62 | 3.48 | 3.56 | **0.01** | 5.07 | 5.85 | **0.00** | 2.14 | 1.64 | 0.07 | 1.95 | 1.36 | 0.09 |
| +5°C | 1.11 | 0.16 | 0.30 | 2.98 | 2.85 | *0.02* | 2.25 | 1.80 | 0.06 | 3.28 | 3.28 | *0.02* | 3.27 | 3.26 | *0.02* |

**Table B13.** Results of Anderson-Darling tests comparing median non-N-fixing cyanobacterial maximum biomass per year between the uniform distribution and each of the variable distributions (Normal, Poisson- 2, 4, or 8 Tail, denoted by Distrib.) for each air temperature offset. The Anderson-Darling criterion (AD), standardized test statistic (T.AD), and *P*-values (asymptotic approximation) are reported for each pairwise scenario comparison. *P*-values in bold indicate significant (Bonferroni-corrected α = 0.01) shifts in the distribution; values in italics indicate shifts with marginal significance (Bonferroni-corrected α = 0.02).

| **Distrib.** | | **+0** | | | **+1** | | | **+2** | | | **+3** | | | **+4** | | | **+5** | |
|---|---|---|---|---|---|---|---|---|---|---|---|---|---|---|---|---|---|---|
| | **AD** | **T.AD** | ***P*** | **AD** | **T.AD** | ***P*** | **AD** | **T.AD** | ***P*** | **AD** | **T.AD** | ***P*** | **AD** | **T.AD** | ***P*** | **AD** | **T.AD** | ***P*** |
| Normal | 0.41 | -0.84 | 0.89 | 1.64 | 0.91 | 0.14 | 0.54 | -0.66 | 0.75 | 0.57 | -0.61 | 0.71 | 0.35 | -0.94 | 0.95 | 0.36 | -0.91 | 0.41 |
| Poisson-Tail 2 | 0.00 | -1.44 | 1.00 | 1.54 | 0.78 | 0.16 | 0.50 | -0.72 | 0.79 | 1.74 | 1.07 | 0.12 | 0.25 | -1.07 | 1.00 | 0.47 | -0.76 | **0.00** |
| Poisson-Tail 4 | 0.00 | -1.44 | 1.00 | 0.83 | -0.24 | 0.47 | 0.59 | -0.59 | 0.70 | 1.47 | 0.68 | 0.17 | 0.36 | -0.92 | 0.94 | 0.48 | -0.75 | **0.00** |
| Poisson-Tail 8 | 0.00 | -1.44 | 1.00 | 0.92 | -0.12 | 0.41 | 0.80 | -0.28 | 0.49 | 1.14 | 0.20 | 0.29 | 0.28 | -1.04 | 0.99 | 0.65 | -0.51 | **0.00** |

**Table B14.** Results of Anderson-Darling tests comparing median N-fixing cyanobacterial maximum biomass per year between the uniform distribution and each of the variable distributions (Normal, Poisson- 2, 4, or 8 Tail, denoted by Distrib.) for each air temperature offset. The Anderson-Darling criterion (AD), standardized test statistic (T.AD), and *P*-values (asymptotic approximation) are reported for each pairwise scenario comparison. *P*-values in bold indicate significant (Bonferroni-corrected α = 0.01) shifts in the distribution; values in italics indicate shifts with marginal significance (Bonferroni-corrected α = 0.02).

| **Distribution** | | **+0** | | | **+1** | | | **+2** | | | **+3** | | | **+4** | | | **+5** | |
|---|---|---|---|---|---|---|---|---|---|---|---|---|---|---|---|---|---|---|
| | **AD** | **T.AD** | ***P*** | **AD** | **T.AD** | ***P*** | **AD** | **T.AD** | ***P*** | **AD** | **T.AD** | ***P*** | **AD** | **T.AD** | ***P*** | **AD** | **T.AD** | ***P*** |
| Normal | 0.61 | -0.57 | 0.68 | 0.43 | -0.82 | 0.87 | 0.55 | -0.65 | 0.74 | 0.52 | -0.69 | 0.77 | 0.51 | -0.70 | 0.78 | 0.26 | -1.07 | 0.99 |
| Poisson-Tail 2 | 0.00 | -1.44 | 1.00 | 0.79 | -0.30 | 0.50 | 0.41 | -0.84 | 0.89 | 0.52 | -0.69 | 0.77 | 0.61 | -0.56 | 0.67 | 0.26 | -1.07 | 0.99 |
| Poisson-Tail 4 | 0.00 | -1.44 | 1.00 | 0.74 | -0.38 | 0.55 | 0.49 | -0.74 | 0.81 | 0.37 | -0.91 | 0.93 | 0.43 | -0.82 | 0.87 | 0.26 | -1.07 | 0.99 |
| Poisson-Tail 8 | 0.00 | -1.44 | 1.00 | 0.59 | -0.59 | 0.70 | 0.62 | -0.54 | 0.66 | 0.52 | -0.69 | 0.77 | 0.46 | -0.77 | 0.84 | 0.26 | -1.07 | 0.99 |

**Table B15.** Results of Anderson-Darling tests comparing median non-N-fixing cyanobacterial maximum biomass per year between the baseline scenario (+0°C) and each of the warming distribution scenarios across air temperature offsets (+1°C to +5°C). The Anderson-Darling criterion (AD), standardized test statistic (T.AD), and *P*-values (asymptotic approximation) are reported for each pairwise scenario comparison. *P*-values in bold indicate significant (Bonferroni-corrected α = 0.01) shifts in the distribution; values in italics indicate shifts with marginal significance (Bonferroni-corrected α = 0.02).

| Temp. Offset | Uniform | | | Normal | | | Poisson- Tail 2 | | | Poisson- Tail 4 | | | Poisson- Tail 8 | | |
|---|---|---|---|---|---|---|---|---|---|---|---|---|---|---|---|
| | AD | T.AD | *P* | AD | T. AD | *P* | AD | T. AD | *P* | AD | T. AD | *P* | AD | T. AD | *P* |
| +1°C | 3.06 | 2.95 | *0.02* | 1.30 | 0.42 | 0.23 | 1.52 | 0.75 | 0.16 | 2.08 | 1.55 | 0.07 | 1.64 | 0.92 | 0.13 |
| +2°C | 0.94 | -0.09 | 0.39 | 2.07 | 1.54 | 0.07 | 1.82 | 1.17 | 0.11 | 1.88 | 1.26 | 0.10 | 1.88 | 1.26 | 0.10 |
| +3°C | 0.26 | -1.07 | 0.99 | 0.71 | -0.41 | 0.57 | 2.12 | 1.61 | 0.07 | 1.66 | 0.95 | 0.13 | 1.07 | 0.10 | 0.32 |
| +4°C | 1.44 | 0.63 | 0.18 | 0.80 | -0.29 | 0.49 | 1.52 | 0.75 | 0.16 | 1.21 | 0.30 | 0.26 | 1.57 | 0.82 | 0.15 |
| +5°C | 0.59 | -0.59 | 0.70 | 0.36 | -0.91 | 0.94 | 0.75 | -0.36 | 0.54 | 0.45 | -0.79 | 0.85 | 0.21 | -1.13 | 1.00 |

**Table B16.** Results of Anderson-Darling tests comparing median N-fixing cyanobacterial maximum biomass per year between the baseline scenario (+0°C) and each of the warming distribution scenarios across air temperature offsets (+1°C to +5°C). The Anderson-Darling criterion (AD), standardized test statistic (T.AD), and *P*-values (asymptotic approximation) are reported for each pairwise scenario comparison. *P*-values in bold indicate significant (Bonferroni-corrected α = 0.01) shifts in the distribution; values in italics indicate shifts with marginal significance (Bonferroni-corrected α = 0.02).

| **Temp. Offset** | **Uniform** | | | **Normal** | | | **Poisson- Tail 2** | | | **Poisson- Tail 4** | | | **Poisson- Tail 8** | | |
|---|---|---|---|---|---|---|---|---|---|---|---|---|---|---|---|
| | **AD** | **T.AD** | ***P*** | **AD** | **T. AD** | ***P*** | **AD** | **T. AD** | ***P*** | **AD** | **T. AD** | ***P*** | **AD** | **T. AD** | ***P*** |
| +1°C | 0.70 | -0.44 | 0.59 | 0.74 | -0.37 | 0.54 | 0.63 | -0.53 | 0.65 | 0.68 | -0.46 | 0.60 | 0.44 | -0.81 | 0.86 |
| +2°C | 1.16 | 0.23 | 0.28 | 0.54 | -0.67 | 0.76 | 0.60 | -0.58 | 0.69 | 0.60 | -0.58 | 0.69 | 0.54 | -0.67 | 0.76 |
| +3°C | 0.54 | -0.66 | 0.75 | 0.46 | -0.78 | 0.84 | 0.73 | -0.39 | 0.56 | 0.62 | -0.55 | 0.66 | 0.73 | -0.39 | 0.56 |
| +4°C | 0.95 | -0.07 | 0.39 | 0.65 | -0.51 | 0.64 | 0.48 | -0.75 | 0.82 | 0.84 | -0.23 | 0.46 | 0.42 | -0.83 | 0.88 |
| +5°C | 0.64 | -0.52 | 0.64 | 0.84 | -0.23 | 0.46 | 0.95 | -0.08 | 0.39 | 0.78 | -0.32 | 0.51 | 0.92 | -0.11 | 0.40 |

**Table B17.** Results of Anderson-Darling tests comparing maximum non-N-fixing maximum biomass per year between the baseline scenario (+0°C) and each of the warming distribution scenarios across air temperature offsets (+1°C to +5°C). The Anderson-Darling criterion (AD), standardized test statistic (T.AD), and *P*-values (asymptotic approximation) are reported for each pairwise scenario comparison. *P*-values in bold indicate significant (Bonferroni-corrected α = 0.01) shifts in the distribution; values in italics indicate shifts with marginal significance (Bonferroni-corrected α = 0.02).

| Temp. Offset | Uniform | | | Normal | | | Poisson- Tail 2 | | | Poisson- Tail 4 | | | Poisson- Tail 8 | | |
|---|---|---|---|---|---|---|---|---|---|---|---|---|---|---|---|
| | AD | T.AD | *P* | AD | T. AD | *P* | AD | T. AD | *P* | AD | T. AD | *P* | AD | T. AD | *P* |
| +1°C | 3.06 | 2.95 | *0.02* | 5.38 | 6.28 | **0.00** | 6.51 | 7.92 | **0.00** | 7.75 | 9.70 | **0.00** | 6.45 | 7.83 | **0.00** |
| +2°C | 0.94 | -0.09 | 0.39 | 7.07 | 8.72 | **0.00** | 4.93 | 5.64 | **0.00** | 7.07 | 8.72 | **0.00** | 6.45 | 7.83 | **0.00** |
| +3°C | 0.26 | -1.07 | 0.99 | 3.48 | 3.57 | **0.01** | 5.22 | 6.06 | **0.00** | 6.45 | 7.83 | **0.00** | 4.50 | 5.03 | **0.00** |
| +4°C | 1.44 | 0.63 | 0.18 | 3.75 | 3.95 | **0.01** | 6.51 | 7.92 | **0.00** | 3.50 | 3.60 | **0.01** | 4.35 | 4.81 | **0.00** |
| +5°C | 0.59 | -0.59 | 0.70 | 2.85 | 2.66 | 0.03 | 2.99 | 2.86 | *0.02* | 1.92 | 1.33 | 0.09 | 2.33 | 1.91 | 0.05 |

**Table B18.** Results of Anderson-Darling tests for maximum non-N-fixing cyanobacterial maximum biomass per year between the uniform distribution and each of the variable distributions (Normal, Poisson- 2, 4, or 8 Tail, denoted by Distrib.) for each air temperature offset. The Anderson-Darling criterion (AD), standardized test statistic (T.AD), and *P*-values (asymptotic approximation) are reported for each pairwise scenario comparison. *P*-values in bold indicate significant (Bonferroni-corrected α = 0.01) shifts in the distribution; values in italics indicate shifts with marginal significance (Bonferroni-corrected α = 0.02).

| **Distribution** | | **+0** | | | **+1** | | | **+2** | | | **+3** | | | **+4** | | | **+5** | |
|---|---|---|---|---|---|---|---|---|---|---|---|---|---|---|---|---|---|---|
| | **AD** | **T.AD** | ***P*** | **AD** | **T.AD** | ***P*** | **AD** | **T.AD** | ***P*** | **AD** | **T.AD** | ***P*** | **AD** | **T.AD** | ***P*** | **AD** | **T.AD** | ***P*** |
| Normal | 1.74 | 1.06 | 0.12 | 0.60 | -0.58 | 0.69 | 4.07 | 4.41 | **0.01** | 3.23 | 3.21 | *0.02* | 1.14 | 0.21 | 0.28 | 2.12 | 1.61 | 0.07 |
| Poisson-Tail 2 | 0.00 | -1.44 | 1.00 | 1.70 | 1.00 | 0.12 | 2.40 | 2.01 | 0.05 | 5.14 | 5.94 | **0.00** | 2.68 | 2.41 | 0.03 | 2.71 | 2.46 | 0.03 |
| Poisson-Tail 4 | 0.00 | -1.44 | 1.00 | 2.22 | 1.75 | 0.06 | 3.75 | 3.96 | **0.01** | 5.91 | 7.05 | **0.00** | 0.77 | -0.33 | 0.52 | 1.30 | 0.42 | 0.23 |
| Poisson-Tail 8 | 0.00 | -1.44 | 1.00 | 1.49 | 0.71 | 0.17 | 3.32 | 3.33 | *0.02* | 3.04 | 2.93 | *0.02* | 1.19 | 0.27 | 0.27 | 1.86 | 1.24 | 0.10 |

**Table B19.** Results of Anderson-Darling tests comparing maximum N-fixing cyanobacterial maximum biomass per year between the baseline scenario (+0°C) and each of the warming distribution scenarios across air temperature offsets (+1°C to +5°C). The Anderson-Darling criterion (AD), standardized test statistic (T.AD), and *P*-values (asymptotic approximation) are reported for each pairwise scenario comparison. *P*-values in bold indicate significant (Bonferroni-corrected α = 0.01) shifts in the distribution; values in italics indicate shifts with marginal significance (Bonferroni-corrected α = 0.02).

| Temp. Offset | Uniform | | | Normal | | | Poisson- Tail 2 | | | Poisson- Tail 4 | | | Poisson- Tail 8 | | |
|---|---|---|---|---|---|---|---|---|---|---|---|---|---|---|---|
| | AD | T.AD | *P* | AD | T. AD | *P* | AD | T. AD | *P* | AD | T. AD | *P* | AD | T. AD | *P* |
| +1°C | 0.70 | -0.44 | 0.59 | 4.23 | 4.65 | **0.01** | 4.29 | 4.73 | **0.00** | 2.63 | 2.34 | 0.04 | 2.75 | 2.51 | 0.03 |
| +2°C | 1.16 | 0.23 | 0.28 | 2.66 | 2.39 | 0.03 | 2.70 | 2.44 | 0.03 | 2.11 | 1.60 | 0.07 | 3.57 | 3.69 | **0.01** |
| +3°C | 0.54 | -0.66 | 0.75 | 1.93 | 1.33 | 0.09 | 2.79 | 2.57 | 0.03 | 2.56 | 2.24 | 0.04 | 1.68 | 0.97 | 0.13 |
| +4°C | 0.95 | -0.07 | 0.39 | 1.58 | 0.84 | 0.15 | 2.81 | 2.60 | 0.03 | 1.23 | 0.32 | 0.25 | 1.51 | 0.73 | 0.16 |
| +5°C | 0.64 | -0.52 | 0.64 | 1.82 | 1.18 | 0.11 | 1.28 | 0.40 | 0.23 | 1.84 | 1.21 | 0.10 | 3.12 | 3.04 | *0.02* |

**Table B20.** Results of Anderson-Darling tests for maximum N-fixing cyanobacterial maximum biomass per year between the uniform distribution and each of the variable distributions (Normal, Poisson- 2, 4, or 8 Tail, denoted by Distrib.) for each air temperature offset. The Anderson-Darling criterion (AD), standardized test statistic (T.AD), and *P*-values (asymptotic approximation) are reported for each pairwise scenario comparison. *P*-values in bold indicate significant (Bonferroni-corrected α = 0.01) shifts in the distribution; values in italics indicate shifts with marginal significance (Bonferroni-corrected α = 0.02).

| **Distribution** | | **+0** | | | **+1** | | | **+2** | | | **+3** | | | **+4** | | | **+5** | |
|---|---|---|---|---|---|---|---|---|---|---|---|---|---|---|---|---|---|---|
| | **AD** | **T.AD** | ***P*** | **AD** | **T.AD** | ***P*** | **AD** | **T.AD** | ***P*** | **AD** | **T.AD** | ***P*** | **AD** | **T.AD** | ***P*** | **AD** | **T.AD** | ***P*** |
| Normal | 3.92 | 4.20 | **0.01** | 2.75 | 2.51 | 0.03 | 4.58 | 5.14 | **0.00** | 1.37 | 0.53 | 0.20 | 2.54 | 2.21 | 0.04 | 2.44 | 2.06 | 0.05 |
| Poisson-Tail 2 | 0.00 | -1.44 | 1.00 | 2.53 | 2.19 | 0.04 | 4.87 | 5.56 | **0.00** | 1.74 | 1.07 | 0.12 | 4.45 | 4.96 | **0.00** | 1.64 | 0.92 | 0.14 |
| Poisson-Tail 4 | 0.00 | -1.44 | 1.00 | 1.77 | 1.11 | 0.11 | 3.44 | 3.51 | **0.01** | 2.17 | 1.69 | 0.07 | 2.29 | 1.85 | 0.06 | 2.48 | 2.13 | 0.04 |
| Poisson-Tail 8 | 0.00 | -1.44 | 1.00 | 1.91 | 1.31 | 0.09 | 5.11 | 5.91 | **0.00** | 1.10 | 0.14 | 0.30 | 1.92 | 1.32 | 0.09 | 2.93 | 2.78 | *0.02* |
